# DeepSCENIC: transfer learning from sequence-to-function models enables causal gene regulatory network inference

**DOI:** 10.64898/2026.09.18.752607

**Authors:** Gabriele Partel, Seppe De Winter, Vasileios Konstantakos, Casper H. Blaauw, Stein Aerts

## Abstract

Sequence-to-function (S2F) deep learning models have become an important aid to decipher the genomic *cis-*regulatory code. However, current S2F models do not take the cellular *trans-*environment of transcription factors (TF) into account. Conversely, methods for gene regulatory network (GRN) inference often rely on heuristics or simple position weight matrices (PWMs), without exploiting the combinatorial grammar of genomic enhancers. Here, we present DeepSCENIC, a deep learning framework that enables causal GRN inference by performing transfer learning from S2F models to single-cell multiome atlases. We first test and validate DeepSCENIC on ENCODE cell lines, demonstrating that the framework accurately predicts single-cell gene expression and chromatin accessibility by leveraging pretrained S2F models like Enformer and Borzoi. We show that DeepSCENIC recovers TF-region (TF-RE) interactions with high fidelity, and captures de novo TF binding motifs without prior PWM knowledge. The model improves enhancer-gene associations (RE-TG) over correlation-based baselines when benchmarked against large-scale CRISPRi screens. After training a DeepSCENIC model, it enables the prediction of perturbation effects during cell state changes by acting as a mechanistic simulator. In a melanoma cell line atlas, the model accurately recapitulates the transcriptional shift from melanocytic to mesenchymal states, and predicted knock-down effects show high concordance with experimental time series data. Finally, we use DeepSCENIC to identify mouse-human cortex conserved GRNs, finding high cross-species concordance in TF activity programs across matched neuronal subclasses and validating top-ranked enhancers against experimental reporter assays. By unifying S2F enhancer representations with single-cell multiomics, DeepSCENIC provides a new paradigm for jointly modeling and simulating *cis-*sequence and *trans-*cellular perturbations.

## Introduction

Sequence-to-function (S2F) deep learning models have revolutionized our understanding of the *cis-*regulatory code that governs the development and function of different cell types in an organism. A first category of S2F models represents (dilated) convolutional networks trained on chromatin accessibility or transcription factor (TF) binding data^1–6^. Such models use a small receptive field (e.g., 2 kilobase input DNA sequences), and biological discoveries are based on saliency (e.g., integrated gradient^7^, DeepLIFT^8^), pattern discovery^9^, and in silico mutagenesis. They have been successfully used to decipher the sequence logic of cell type specific enhancers^10–12^, to compare enhancers across species^13–15^, to design synthetic enhancers^16^, and to extract combinatorial rules of TF binding sites^17^. A second category of S2F models combines convolutional with transformer layers, uses a larger receptive (e.g., 500 kilobases or 1 megabase), and predicts both chromatin features and gene expression^18–20^. A key advantage of modeling the sequence of entire gene loci is that, in principle, enhancer-to-target gene associations can be learned through attention, although the precision and recall of such predictions needs further improvement, and is usually limited to distances to the transcription start site of around ∼50 kilobases^21^. Biological discovery with these models has been mostly focused on the prediction of the effect of sequence variants^22,23^. A third category of models are masked genomic language models^24–27^, trained in a self-supervised manner on large collections of genomes. Biological discoveries with such models on the genomic regulatory code have been more limited, although sequence features related to promoters, splicing and translational regulation have been identified in bacteria^28^ and fungi^29^.

A key limitation of S2F models, besides the challenge of enhancer-gene links and cell type specificity, is that they are exclusively focused on *cis-*regulatory sequences, and do not take the *trans-*environment into account. In other words, even though candidate TF binding sites can be discovered with high confidence, S2F models have no notion of the identity of the TF proteins that recognize and bind to these sites. Linking candidate TFs to model-derived sequence features is currently done by downstream, manual efforts that are based on the correlation of TF expression and motif occurrence^5^. Existing methods that infer gene regulatory networks (GRN), with regulatory connections between TF proteins, their genomic binding sites, and their putative target genes, do not exploit the rich representation of regulatory sequences from S2F models. Rather, state of the art GRN inference methods, such as SCENIC+^30^, Pando^31^, and CellOracle^32^, predict GRNs from single-cell multiome data through heuristics, statistical enrichment, and TF binding site predictions through position weight matrices (PWM). Alternatively, recent deep learning–based GRN inference methods trained on single-cell multiome atlases^33,34^ incorporate chromatin accessibility and TF– gene associations, yet they do not exploit pretrained sequence-to-function models to derive TF–region interactions directly from DNA sequence, nor do they integrate nucleotide-level *cis-*regulatory grammar into the GRN model. Finally, current GRN inference methods focus mostly on static GRNs, with some efforts on the prediction of TF perturbations^30,32^. For example, SCENIC+ simulates cell state changes through GRN perturbations by training random forest regression models on the static GRN topology.

Recently, several studies have begun to bridge S2F models with cellular *trans-*environment. For example, Corgi^35^ introduces a context-aware S2F model that integrates long-range DNA sequence representations with quantitative expression levels of transcription factors. Similarly, Scooby^36^ combines sequence-derived features with single-cell multiomics embeddings, enabling context-dependent regulatory prediction in individual cells. These approaches represent an important step toward combining *cis-*regulatory sequence modeling with the *trans-*regulatory cellular state. However, they do not explicitly parameterize gene regulatory networks as interpretable TF–region–target gene graphs, nor are they designed as mechanistic simulators that propagate *trans-*regulatory perturbations through a learned *cis-*regulatory architecture. As a result, they remain primarily predictive rather than explicitly causal in their formulation.

Here we develop DeepSCENIC, a deep learning framework that can infer GRNs from single cell multiome atlases, through transfer learning from S2F models. Rather than using S2F models purely for variant effect prediction or for extracting rules of *cis*-regulation, we repurpose their learned *cis*-regulatory embeddings to explicitly parameterize TF–region interactions at nucleotide resolution. By integrating these sequence-derived enhancer representations with single-cell multiome measurements, we jointly infer TF–enhancer and enhancer–gene connections, yielding an explicit, TF–region–target gene network. This design allows DeepSCENIC to move beyond context-aware prediction toward a structured regulatory architecture in which *trans*-regulatory perturbations can be propagated through learned *cis*-regulatory links and target genes.

## RESULTS

### DeepSCENIC: Extracting gene regulatory networks from sequence-to-function models

DeepSCENIC is a deep learning framework designed for inferring GRNs by finetuning S2F models or DNA language models with single-cell RNA and ATAC multiome atlases (Fig. 1 and Methods). Specifically, a S2F model is used to infer sequence representations of scATAC-seq peaks, from which binding potentials are predicted for each TF. The binding potential of each TF is then combined with its expression vector from the scRNA-seq data (Fig. 1a). TF activities on each enhancer are expressed as a linear combination of TF expression and its sequence-derived binding potential. Enhancer activities of each ATAC peak are subsequently expressed as the total contribution of TF activities on the enhancer. Finally, the expression of target genes is predicted as the weighted additive contribution of enhancer activities within 1Mb up-and down-stream of its transcription start site (Fig. 1b).

**Figure 1.**
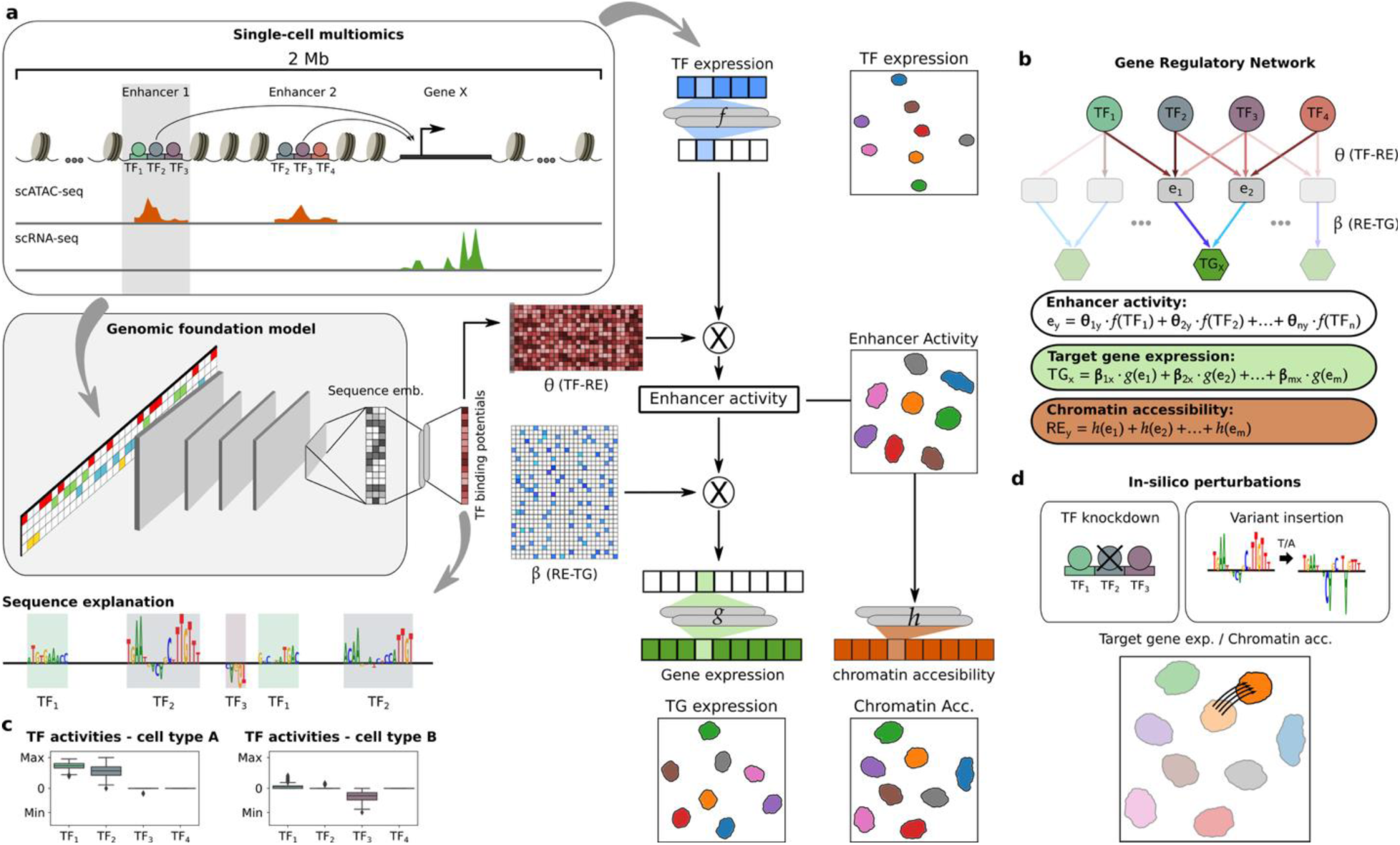
DeepSCENIC model architecture. **(a)** DeepSCENIC leverages S2F models fine-tuned with single-cell multiome data (scATAC-seq and scRNA-seq) to learn regulatory interactions at single-cell level. The S2F model generates sequence embeddings from consensus peak DNA sequences, predicting binding potentials for transcription factors (TFs). TF binding potentials are combined with TF expression to calculate enhancer (regulatory element, RE) activities. Enhancer activities subsequently determine target gene (TG) expression and chromatin accessibility. **(b)** The regulatory interactions between TF to RE, RE to TG (*cis-*regulation), and TF to TG (*trans-* regulation), are learned during model training, resulting in a GRN with single-cell resolution, where the strength of regulatory interactions is captured by parameters (θ and β) learned directly from the data during training. **(c)** DeepSCENIC provides nucleotide-level sequence explanations for TF binding activities on each enhancer, showing differences in TF activities across distinct cell types. **(d)** Post-training, DeepSCENIC enables in-silico perturbations, such as TF knockdowns or sequence variant insertions, assessing their impacts on gene expression and chromatin accessibility by directly modeling perturbed regulatory interactions.

The inferred regulatory interactions between TF-region (TF-RE), region-target gene (RE-TG) and TF-target gene (TF-TG) are learned during training by finetuning a chosen S2F model, optimizing the reconstruction of the target gene expression and chromatin accessibility in each single cell. Thus, the inputs of the model are TF expression vectors of a single cell, and one-hot-encoded DNA sequences. After training, DeepSCENIC yields a GRN at single-cell resolution, reporting TF activities on each scATAC-seq peak with nucleotide-level resolution (Fig. 1c), as well as enhancer to target gene regulatory effects. Crucially, the resulting DeepSCENIC model can be used for in-silico perturbation by propagating the effect of TF expression changes (e.g. TF knockdown simulations) or sequence variants in the input sequences (Fig. 1d and Methods). The result of the perturbation leads to changes in gene expression and chromatin accessibility, which can be mechanistically interpreted using the model’s learned regulatory interactions.

### Validation of DeepSCENIC GRN inference on ENCODE cell-lines

To benchmark the accuracy of the GRNs inferred by DeepSCENIC we trained a model on eight ENCODE deeply profiled cell lines (Fig. 2a, Methods). We evaluated five DeepSCENIC models, each using different S2F or language models: two S2F models that were pretrained on human and mouse ENCODE genomic tracks (i.e. *dS-Enformer* using Enformer^18^ and *dS-Borzoi* using Borzoi^19^); a genomic language model pretrained on a human reference genome (i.e. *dS-HyenaDNA* using HyenaDNA tiny-1k model^24^); a CREsted model^5^ pretrained on chromatin accessibility topics (i.e. *dS-Crested*), and a baseline model (i.e. *dS-CNN)* using a CREsted convolutional neural network trained from scratch (Methods). We evaluated the predictive performance of the five models, by measuring their ability to predict gene expression and chromatin accessibility from TF expression profiles and genomic sequences (Fig. 2b). Performance was assessed on held-out genes and sequences. Among the tested architectures, models leveraging pretrained S2F models (dS-Enformer and dS-Borzoi) achieved the best performance, with dS-Enformer reaching a mean correlation of 0.41 for expression and 0.46 for accessibility. Models without pretraining (e.g. dS-CNN) showed lower performance, underscoring the benefit of transfer learning from large pre-trained S2F models.

**Figure 2.**
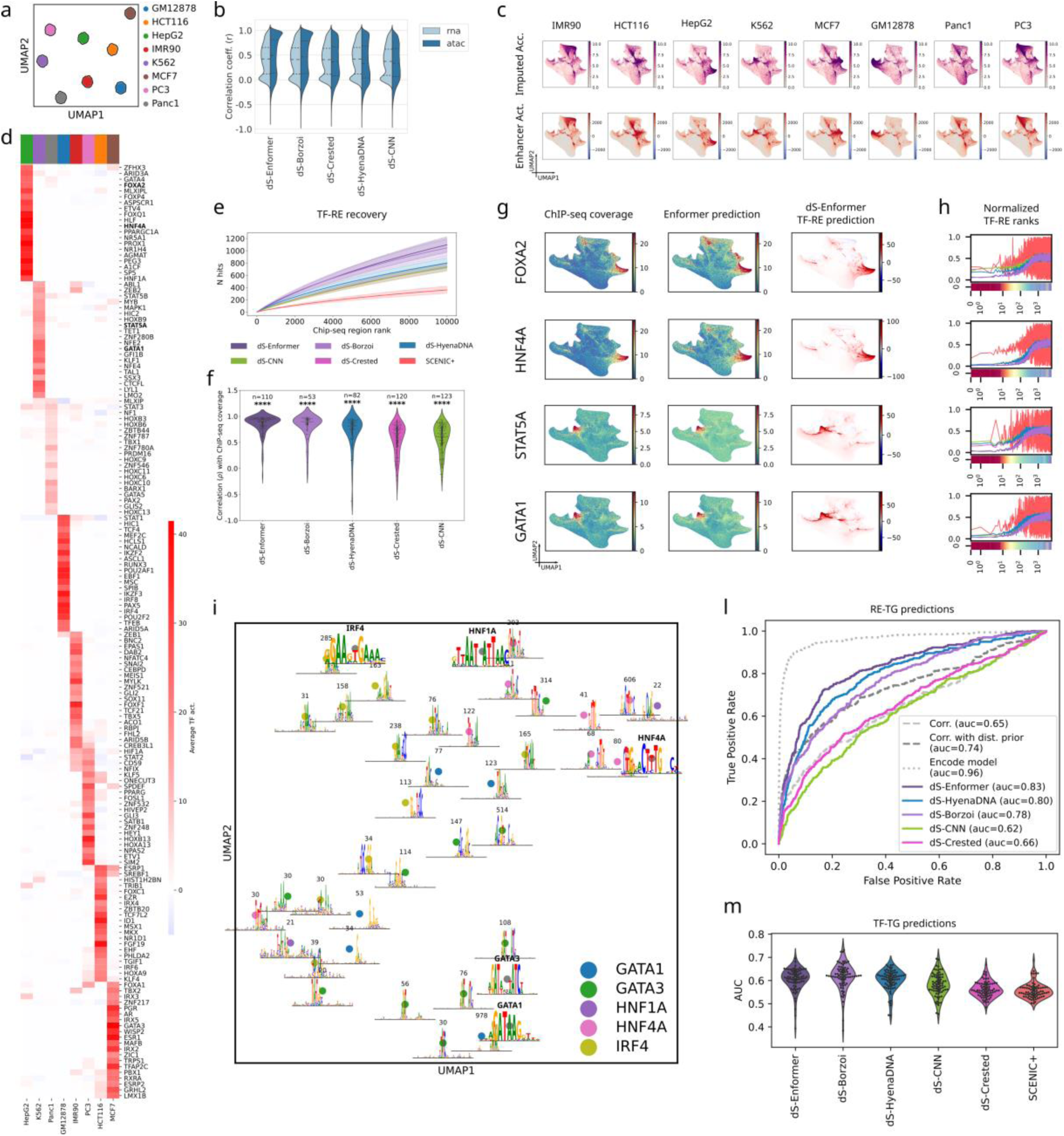
Validation of DeepSCENIC regulatory interactions in ENCODE cell lines. **(a)** UMAP projection of simulated single-cell multiome profiles derived from eight ENCODE cell lines. **(b)** Reconstruction performance on held-out data. Violin plots show Pearson correlation between predicted and observed gene expression (RNA) and chromatin accessibility (ATAC) across model variants using different S2F -backbones. **(c)** UMAP projections of sequence-derived enhancer representations. Top row: chromatin accessibility (imputed scATAC-seq) per cell line. Bottom row: DeepSCENIC-predicted enhancer activities. **(d)** Heatmap of top cell-line–specific transcription factors ranked by average predicted activity over differentially active enhancers. **(e)** TF–RE recovery curves. Cumulative recall of ENCODE ChIP-seq peaks among top-10k ranked predicted TF-bound regions. **(f)** Distribution of per-TF Pearson correlations between predicted TF–RE binding potentials and ENCODE ChIP-seq coverage. Each point represents one TF passing method-specific filtering and with available ChIP-seq data (*n* indicates the number of TFs per method). Significance was computed using a Wilcoxon rank-sum test comparing observed TF–RE correlations to correlations obtained from permuted controls, followed by Holm correction for multiple testing. Asterisks denote adjusted *P* values (*ns* ≥ 0.05, *\** < 0.05, ** < 0.01, *** < 0.001, **** < 0.0001). **(g)** Representative TF binding landscapes for FOXA2, HNF4A, STAT5A, and GATA1. ENCODE ChIP-seq coverage (left), Enformer-predicted signal (middle), and DeepSCENIC TF–RE binding potentials (right) are projected onto the enhancer embedding UMAP. **(h)** Genomic regions were ordered by ChIP-seq signal for representative TFs (GATA1, HNF4A, FOXA2 and STAT5A) and grouped into bins of 100 regions. Line plots show the mean normalized rank assigned by each method within each ChIP-seq bin. The heatmap below each plot shows the corresponding ChIP-seq signal across bins. **(i)** TF motif recovery from in silico sequence evolution. Seqlets extracted from DeepSCENIC-evolved sequences for specific TFs (i.e. GATA1, GATA3, HNF1A, HNF4A, IRF4) were co-embedded with reference motifs; inset logos depict motif patterns and labels report cluster sizes (number of seqlets per pattern). **(l)** RE-TG validation against CRISPRi data in K562 cells. ROC curves and AUC values are shown for predicted RE–TG scores from each method. **(m)** TF-TG validation using ENCODE TF knockout RNA-seq datasets. Violin plots show per-TF area under the ROC curve (AUC) for predicting differentially expressed genes.

To investigate whether DeepSCENIC captures the regulatory specificity of each cell type, we visualized the sequence-derived representations of candidate enhancers (Fig. 2c). Each enhancer sequence was embedded using the learned binding potential scores of all TFs. The resulting projections revealed clear separation of enhancers by cell type (Fig. 2c), suggesting that the model has learned to distinguish cell-type-specific combinations of TF binding sites. In addition, when visualizing the predicted enhancer activities in the same space, we observed that their spatial distribution closely mirrored the chromatin accessibility profiles of each cell type (Fig. 2c). This colocalization indicates that DeepSCENIC not only captures the cell-type-specific regulatory code, but also that the inferred enhancer activities recapitulate chromatin accessibility.

To identify key regulators driving cell-type-specific enhancer activities, we examined the TFs with the highest average predicted activity over differentially active enhancers for each cell line (Methods). We defined differentially active enhancers as those with high predicted activity relative to other cell types (Wilcoxon rank-sum test, Methods), and ranked transcription factors based on their predicted activity across these regions (Methods). The resulting heatmap highlights cell line-specific TFs (Fig. 2d). For example, HNF4A, HNF1A, FOXA2, and GATA4 are predicted as top regulators in HepG2 (hepatocellular carcinoma-derived cell line). These TFs are well-known hepatic master regulators^37^. In K562 (chronic myelogenous leukemia cell line of erythro-myeloid origin), DeepSCENIC identified STAT5A/B and GATA1 as top regulators, again confirmed TFs for K562^38–40^. Next, we sought to validate the inferred *trans-*(TF– RE, TF–TG) and *cis*-(RE–TG) regulatory interactions using independent genomic datasets.

To evaluate the accuracy of TF–RE predictions, we compared DeepSCENIC-inferred TF binding scores to experimental ChIP-seq data from ENCODE. We computed recovery curves showing the number of ChIP-seq peaks recalled among the top 10,000 peaks ranked by DeepSCENIC (Methods). All five DeepSCENIC models recovered a significant number of ChIP-seq peaks, with stronger enrichment compared to SCENIC+ (Fig. 2e). The pretrained S2F DeepSCENIC models (dS-Enformer and dS-Borzoi) achieve the highest recall rates across most TFs. Complementary to this, we computed Pearson correlations between DeepSCENIC TF–RE scores and ChIP-seq signal intensities across candidate regions (Fig. 2f, Methods). Again, dS-Enformer and dS-Borzoi exhibited the strongest concordance with experimental data, with mean correlation coefficients of 0.85 and 0.82 respectively, against 0.58 for the baseline model (dS-CNN). These results demonstrate that DeepSCENIC can accurately recover TF binding landscapes from sequence and TF expression alone, and that pretraining on cell-line data provides a consistent performance boost across evaluation metrics.

We then visualized sequence-derived representations of ATAC peaks for four master regulators: HNF4A and FOXA2 in HepG2, and GATA1 and STAT5A in K562. The DeepSCENIC TF-RE predictions show strong concordance with experimental ChIP-seq and Enformer predicted ChIP-seq signal (Fig. 2g). Furthermore, rank profiles across regions ordered by ChIP-seq signal revealed that DeepSCENIC variants consistently assign better ranks to experimentally supported binding sites compared to the baseline SCENIC+ (Fig. 2h). Regions with stronger ChIP-seq signal tend to receive higher predicted ranks in DeepSCENIC models, indicating improved prioritization of validated TF binding sites.

DeepSCENIC does not use priors for known TF recognition sequences, such as PWM collections. Instead, it relies on S2F sequence embeddings, combined with TF expression, ATAC, and target gene expression, to learn TF activities on genomic enhancers. We therefore examined whether DeepSCENIC learned TF-specific PWMs de novo. To this end, we performed sequence evolution experiments for five TFs (i.e. GATA1, GATA3, HNF4A, HNF1A, and IRF4) (Methods). Starting from random DNA, we iteratively optimized 1,000 sequences per TF to maximize the specificity of TF–RE scores. From the evolved sequences we extracted high-contributing subsequences (seqlets), which were aggregated into sequence patterns. Motif similarity between patterns was quantified using TOMTOM^41^ similarity scores against reference PWMs, which served as a feature space to compute distances between patterns prior to dimensionality reduction (Fig. 2i) ^42^. For GATA1, GATA3, HNF4A, and IRF4, evolved sequences consistently converged to the expected canonical motif or closely related variants, demonstrating that DeepSCENIC accurately captures their binding specificities. In contrast, sequence optimization for HNF1A failed to converge in most cases, and the minority of sequences that did converge lacked well-formed HNF1A motifs, often resembling HNF4A motifs instead (Fig. 2i). These TFs belong to distinct families with different DNA-binding domains (HNF4A is a nuclear receptor, whereas HNF1A contains a homeodomain-like DNA-binding domain). This reduced specificity likely arises from the extensive co-binding and cooperative regulation of HNF1A and HNF4A in HepG2 cells, where they frequently occupy the same regulatory elements. Given the resolution of the training data (consensus peaks and cell-type expression), the model may lack sufficient discriminative information to fully disentangle their contributions. These findings illustrate the capacity of DeepSCENIC to *de novo* recover high-fidelity sequence motifs for TFs.

Next, we assessed the quality of enhancer-to-target gene (RE–TG) interactions using an experimental benchmark based on a recent CRISPRi screen^43^, in which 10,411 region-gene pairs were profiled in the K562 cell line. For each DeepSCENIC model, we computed receiver operating characteristic (ROC) curves based on predicted RE–TG scores, using experimentally validated links as positives and unlinked pairs as negatives, where validated links correspond to enhancer–gene pairs showing significant expression changes upon CRISPRi perturbation (Benjamini–Hochberg FDR < 0.05) (Fig. 2l, Methods). We compared model performances against two correlation-based baselines: a raw Pearson correlation between enhancer and gene expression, and a distance-aware variant where correlation scores are weighted by a prior based on genomic distance. While the best-performing DeepSCENIC model (dS-Enformer) outperformed both correlation-based baselines, models such as dS-CNN and dS-Crested performed worse or on par with them. All DeepSCENIC models, however, remained inferior to the gold standard supervised model that was trained directly on the CRISPRi data (ENCODE model, AUC = 0.96). dS-Enformer achieved the highest performance among the DeepSCENIC variants, with an AUC of 0.83, dS-HyenaDNA and dS-Borzoi followed with AUCs of 0.80 and 0.78, respectively.

Finally, we computed TF-TG interactions from predicted TF-RE and RE-TG interactions and validated these using TF knock-down experiments. Particularly, we obtained sets of differentially expressed genes for 63 TF knock-down experiments in ENCODE cell lines, and computed the area under the ROC curve (AUC) against DeepSCENIC-predicted TF–TG scores (Fig. 2m, Methods). All DeepSCENIC variants outperformed SCENIC+, with the largest performance gains observed for models leveraging foundation model pretraining. dS-Enformer and dS-Borzoi achieved the highest AUCs (average AUCs 0.6 and 0.61 respectively), followed by dS-HyenaDNA, dS-CNN, dS-Crested and SCENIC+.

These combined validation results illustrate the capacity of integrating both sequence and multiomic data through end-to-end learning, enabling the model to capture context-specific regulatory interactions that are not explained by proximity or expression correlation alone.

### DeepSCENIC simulates cell state changes upon TF and sequence perturbation

To evaluate the utility of DeepSCENIC to model cell state changes upon TF and sequence perturbations, we trained a GRN model on a single-cell multiomic dataset of melanoma cell lines^30^. This dataset comprises 9 melanoma cultures spanning distinct phenotypes, including the melanocytic (MEL), mesenchymal-like (MES), and intermediate (INT) states. A principal component analysis (PCA) of gene expression reveals a continuum of cell states, with PC1 capturing the transition from MEL to MES phenotypes (Fig. 3a). We first inferred state-specific TFs using DeepSCENIC and identified known lineage-determining factors among the top-scoring TFs, including MITF, TFAP2A and SOX10 in MEL states, ETV4 and ELF1 for INT states and NR2F2, TBX3, and NFIX in MES-like cultures (Fig. 3b), consistent with prior reports linking these TFs to MEL-MES identities and phenotype switching in melanoma^44^. Predicted target genes of these TFs include, for the MEL state, pigment-related genes (e.g. MLANA, DCT, PMEL) and proliferation (e.g. DTYMK, RAD51C, ID3, EIF4B) genes downstream of SOX10; and for the MES state target genes linked to stress response, migration, and inflammatory signaling genes (e.g. SMAD3, TNFRSF12A, AXL, RAC2) (Fig 3e).

**Figure 3.**
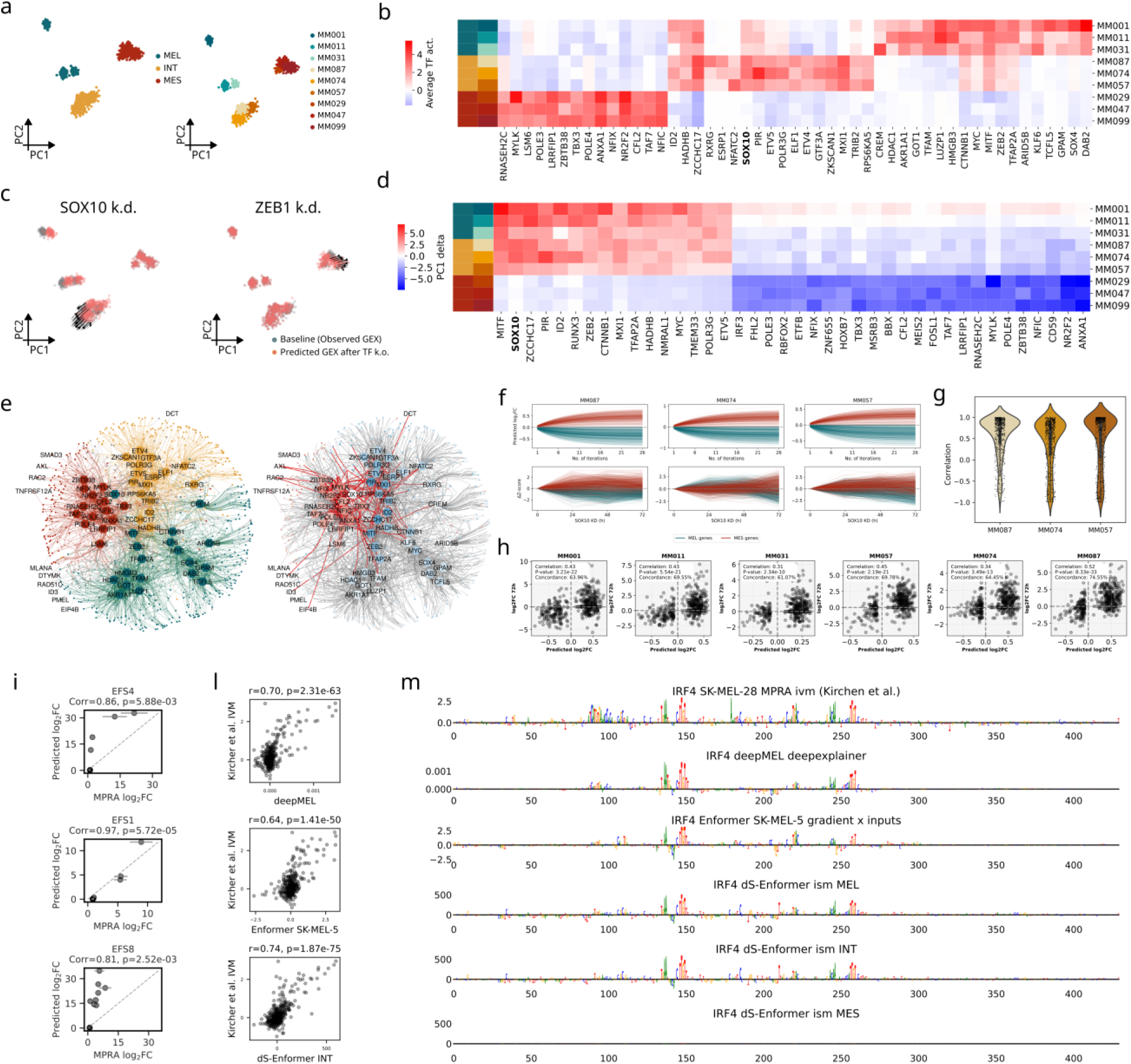
DeepSCENIC simulates cell state transitions and *cis-*regulatory perturbations in melanoma. (a) Principal component analysis (PCA) of gene expression profiles from nine melanoma cell lines spanning melanocytic (MEL), intermediate (INT), and mesenchymal-like (MES) states. Left: cells colored by phenotypic state. Right: cells colored by individual melanoma lines. **(b)** Heatmap of top-ranked transcription factors (TFs) ordered by average predicted TF activity across differentially active enhancers between melanoma lines. **(c)** In silico TF perturbations projected onto the MEL–MES axis (PC1). Predicted gene expression after SOX10 (left) or ZEB1 (right) knockdown is shown relative to baseline. **(d)** Heatmap of predicted shifts along the MEL–MES axis (ΔPC1) following in silico TF knockdown across melanoma lines. Columns show the top 40 TFs ranked by mean absolute ΔPC1 across lines, rows correspond to individual cell lines. **(e)** DeepSCENIC-inferred gene regulatory network in melanoma cell lines. Left: baseline regulatory network colored by phenotypic state (MEL, INT, MES). Right: network colored by predicted regulatory changes following TF perturbation. **(f)** MEL and MES marker gene dynamics following SOX10 knockdown in INT melanoma lines. **Top:** DeepSCENIC-predicted log2 fold-changes of MEL (teal) and MES (red) marker genes across simulation iterations. **Bottom:** experimentally measured changes in marker gene set Δz-scores relative to baseline at 0 h, 24 h, 48 h, and 72 h after SOX10 knockdown for the same marker gene sets. **(g)** Per-gene correlation between predicted and experimental SOX10 knockdown trajectories for curated Verfaillie MEL/MES genes in INT melanoma lines (see Methods). Points represent individual genes; violins show per-line distributions. **(h)** Scatterplots comparing predicted versus experimentally observed log2 fold changes for differentially expressed genes across melanoma lines. Each panel corresponds to one line; Pearson correlation and concordance correlation coefficients are indicated. **(i)** Comparison of DeepSCENIC-predicted mutation effects (log2FC) with experimentally measured MPRA log2FC values for three synthetic melanoma enhancers (EFS4, EFS1, EFS8). Pearson correlation coefficients and P values are indicated. **(l)** Correlation between sequence explanation contribution scores from different models and experimentally measured in vitro mutagenesis (IVM) effects for the IRF4 enhancer. **(m)** Nucleotide-level mutagenesis profiles across the IRF4 enhancer. Shown are experimental MPRA IVM measurements (top), DeepMEL attribution scores, Enformer gradient × input scores, and DeepSCENIC ISM predictions in MEL, INT, and MES cellular contexts.

We next simulated TF knock-downs by ablating individual TF expression and propagating their regulatory effects through the DeepSCENIC-inferred GRN (Methods). Silencing of SOX10 induced a shift along PC1 toward the MES program across MEL-like cell lines (Fig. 3c), consistent with its experimentally validated role as a driver of phenotype switching in melanoma^44^. Consistent with this DeepSCENIC predicts an upregulation of MEL-specific TFs and target genes and a downregulation of MES TFs and target genes (Fig. 3e). By comparison to curated gene signatures associated with MEL and MES phenotypes^45^ (Fig. 3f), the model indeed recapitulates the expected cell state switch toward a mesenchymal phenotype. To further validate these predictions, we compared the predicted changes to the trajectory of expression changes following experimental SOX10 knockdown in three intermediate MEL lines (MM074, MM087, MM057)^44^. MEL and MES program activity scores computed across simulation steps showed progressive depletion of MEL signature genes and concomitant increase in MES gene expression, closely mirroring the transcriptional reprogramming observed experimentally, with strong concordance between predicted log2FC trajectories and experimentally measured changes in standardized MEL/MES program activity scores (Δz-scores relative to baseline) across the SOX10 knockdown time course(Fig. 3f, g). This is also reflected by directly comparing the predicted versus measured log2FC values at the 72-hour time point post SOX10 knock-down for differentially expressed MEL and MES marker genes (adjusted p < 0.05, log2FC > 2). Across MEL and INT lines, we observed significant correlations (Pearson r = 0.31–0.52) and high concordance (Concordance Correlation Coefficient = 0.61–0.75) between predicted and observed expression changes (Fig. 3h). Encouraged by this result, we performed an in silico screen, perturbing each TF in turn, and measuring the predicted effect on the MEL-MES axis. Only a subset of TFs drive transitions, including SOX10, MITF, and RUNX3, whose knock-down are predicted to induce the most pronounced shifts toward the MES state; while perturbations of ANXA1, NR2F2, FOSL1, and ZEB1 led to a predicted inverse transition toward the MEL state (Fig. 3c, d). These results demonstrate that DeepSCENIC can simulate complex transcriptional responses to TF perturbations in a cell-type specific manner, supporting its utility for modeling cell state transitions.

Besides modifying the *trans-*environment, we also investigated whether DeepSCENIC can predict the effect of sequence variants in enhancers. We performed in silico mutagenesis of three MEL-specific synthetic enhancers previously designed and validated in Taskiran, et al.^16^. For each enhancer, we simulated the sequence evolution across multiple steps, progressively introducing mutations, and measured the predicted changes in regulatory activity (log₂FC) using DeepSCENIC. These predictions were then compared to in vitro measurements from luciferase assays^16^, performed on matched mutant sequences. Across all three enhancers, DeepSCENIC predictions showed strong agreement with experimental measurements, with Pearson correlations ranging from 0.81 to 0.97 (Fig. 3i). In another validation experiment, we compared DeepSCENIC in silico mutagenesis (ISM) scores with experimental in vitro mutagenesis (IVM) measurements from Kircher et al.^46^ for the IRF4-enhancer that is active in MEL and INT melanoma states. We benchmarked DeepSCENIC predictions against two other S2F models, namely DeepMEL^14^ and Enformer^18^ (interpreted for the SK-MEL-5 class using gradient × input attributions). Across all models, predicted mutation effects were significantly correlated with IVM scores, with DeepSCENIC achieving the highest concordance (Pearson r = 0.74) compared to DeepMEL (r = 0.70) and Enformer (r = 0.64) (Fig. 3l). Visualization of ISM profiles revealed that DeepSCENIC accurately recovered the core TF binding motifs and captured surrounding sequence context features associated with enhancer activity in MEL and INT states, while predictions for MES states showed minimal signal, consistent with the enhancer’s cell-type specificity (Fig. 3m). These results further support the model’s ability to learn context-dependent enhancer grammar within the nodes of the GRN, and to recapitulate experimentally measured sequence-function relationships.

These results highlight DeepSCENIC ability to model the quantitative effects of sequence variants in a cell-state-specific manner, enabling precise prediction of how single-nucleotide changes or designed mutations alter enhancer function.

### DeepSCENIC identifies conserved GRNs in the human and mouse cerebral cortex

Ground truth, experimentally validated GRNs are virtually non-existent at a genome-wide scale and at cell type specific resolution. While for certain TFs and cell types, genome-wide TF-binding sites are available through ChIP-seq, the functional, causal link to specific target genes remains largely unverified. Even CRISPR-mediated TF knockdowns fail as a definitive ’gold standard’ because the high level of technical noise and the arbitrary choice of time points make it nearly impossible to distinguish immediate direct targets from cascading indirect effects. Likewise, the identification of functionally relevant TFs per cell type lacks ground truth data, besides anecdotal evidence for TF mutants in model organisms that cause clear phenotypes, or human mutations in TFs that cause disease. An alternative approach to assess the validity of predicted TFs per cell type, and predicted target genes per TF, is to rely on comparative genomics. Here, we train DeepSCENIC models on the human and mouse cerebral cortex, using single-cell multiome atlases of the motor cortex (M1) from the BICCN/BICAN consortium^47^ (Fig 5a). Importantly, a validated set of cell type-specific enhancers is available for the cortex cell types identified using AAV enhancer-reporter assays, in the Armamentarium^48^. To assess the conservation of transcription factor (TF) activity programs between mouse and human, we compared inferred TF activities across one-to-one matched cell types (Methods). We first restricted the analysis to one-to-one orthologous TFs with detectable activity in both species. For each matched cortical subclass, TFs were ranked independently in each species according to their inferred activity, and candidate conserved TFs were selected as those present among the top 10% highest-ranked TFs in both species. These candidates were further prioritized using a rank-product criterion, retaining TFs with concordant high ranks across mouse and human. Finally, for each retained TF, conservation of its subclass-specific activity profile was quantified by correlating its activity across matched cortical subclasses between mouse and human, and TFs with correlation greater than 0.5 were retained as conserved regulators (Fig. 4b, Methods). Among these, we find well known regulators as conserved TFs (e.g., MEF2C, CUX2, RORA, RFX3, and MEIS2 for L2/3IT neurons). Quantifying conservation using Jaccard overlap across increasing top-k sets of TFs, defined as the k highest-ranked TFs by inferred activity within each matched cell type and species, revealed substantially higher overlap than expected by chance for all major cortical cell types, with normalized enrichment scores (NES) consistently above random expectation (Fig. 4c, Methods).

**Figure 4.**
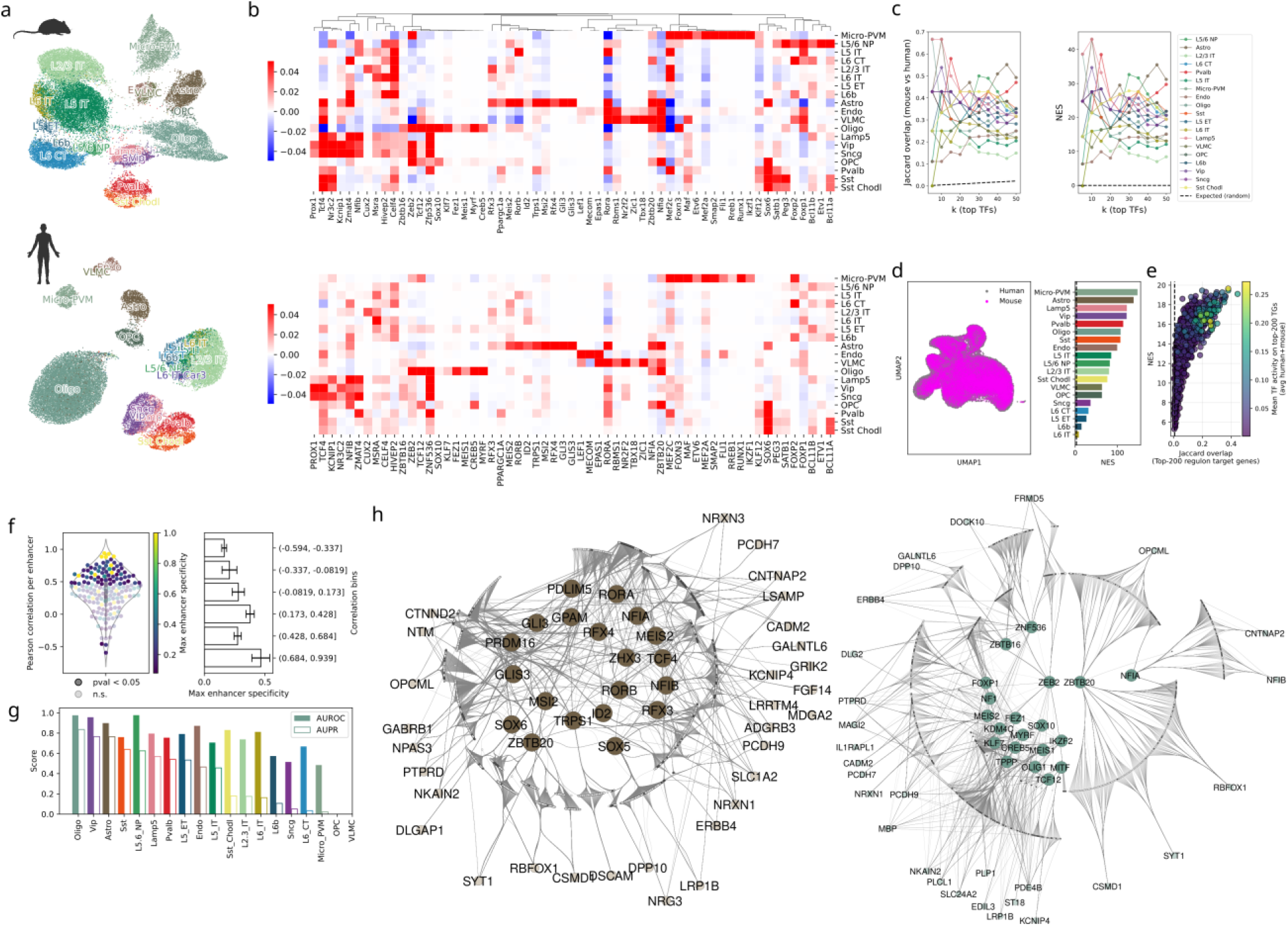
Cross-species conservation of gene regulatory programs in human and mouse motor cortex. **(a)** UMAP projections of DeepSCENIC-inferred enhancer activity profiles in mouse (top) and human (bottom) primary motor cortex (M1), colored by annotated cortical subclasses. **(b)** Heatmaps of DeepSCENIC-inferred TF activity scores across matched cortical subclasses in mouse (top) and human (bottom), restricted to conserved orthologous TFs. **(c)** Cross-species overlap of top-ranked TFs per matched cell type. Left: Jaccard similarity between mouse and human top-k TF sets as a function of k. Right: normalized enrichment scores (NES) relative to random expectation. **(d)** Cross-species enhancer regulatory code conservation. Left: co-embedding of mouse and human enhancers using DeepSCENIC sequence-derived TF binding representations. Right: quantification of neighborhood enrichment, showing the fraction of nearest neighbors belonging to the same annotated cell type from the opposite species, compared to random expectation (see Methods). **(e)** Conservation of TF–TG relationships across species. NES for overlap of top-200 predicted TF target genes per cell type. **(f)** Correlation between predicted enhancer activity and experimentally measured Armamentarium AAV reporter activity. Left: Pearson correlation per enhancer, colored by maximum enhancer specificity. Right: mean enhancer specificity across correlation bins. **(g)** Cell-type–specific enhancer classification performance. Area under the ROC curve (AUROC) and area under the precision–recall curve (AUPR) for predicting experimentally validated enhancer activity per cell type. **(h)** Cross-species conserved gene regulatory networks. Left: astrocyte network showing conserved TF–RE-TG interactions. Right: oligodendrocyte network showing conserved TF–RE-TG interactions. Regulatory element (region) nodes are color-coded based on cross-species conservation after genomic liftover: grey nodes indicate conserved regions, whereas white nodes indicate non-conserved regions.

Having established conservation of TF activity programs, we next asked whether this conservation extends to the underlying *cis-*regulatory sequence code. To this end, we co-embedded mouse and human enhancers using the human model sequence-derived TF-binding predictions, and found that enhancers from the same cell type cluster together regardless of species (Fig. 4d; Supplementary Fig. 1). Neighborhood analysis confirmed that, within each cell type, human enhancers are most similar to mouse enhancers of the same cell type, indicating conservation of the underlying sequence code (Fig. 4d).

To assess conservation at the level of TF–TG gene relationships, we calculated TF-TG links from TF-RE and RE-TG links. Conserved TF–TG relationships were defined by restricting target genes to one-to-one mouse-human orthologs shared among the top–200 predicted targets per TF in both species. While target gene conservation was lower than TF conservation, consistent with known divergence of non-TF gene expression programs between human and mouse^49^, a substantial subset of regulon targets exhibited overlap beyond random expectation, and higher overlap was associated with TFs showing stronger activity in both species (Fig. 4e).

To further contextualize conserved transcriptional regulation within a cell-type–specific framework, we constructed cross-species GRNs for astrocytes and oligodendrocytes by integrating independently inferred mouse and human DeepSCENIC GRNs. For each cell type, GRNs were first reconstructed separately in mouse and human, starting from the shared set of conserved TFs previously defined. Mouse regulatory regions were subsequently lifted over to the human genome and matched to human regulatory regions based on genomic overlap, enabling the identification of conserved enhancer–target associations. The final cross-species GRNs were obtained by merging the mouse and human networks and retaining only TF–TG relationships present in both species, thereby enforcing conservation at the level of transcription factors, regulatory regions, and target genes (Fig. 4h, Methods). The astrocytic network is organized around well-established glial regulators, including RORA, NFIA, NFIB, and SOX6, which form conserved regulatory hubs coordinating astrocyte-specific transcriptional programs. Conserved astrocytic target genes were strongly enriched for synapse-associated processes, including *trans-*synaptic signaling and synapse organization, driven by a coherent adhesion and signaling module (e.g., NRXN1/3, PTPRD, CNTNAP2, ERBB4–NRG3) consistent with astrocyte-mediated regulation of neuron–glia communication.

The oligodendrocytic GRN is dominated by lineage-defining transcription factors such as SOX10, MITF, MYRF, and OLIG family members, forming a densely interconnected regulatory module controlling oligodendrocyte maturation and myelination programs. Under the stringent cross-species intersection used to define conservation, enriched terms among oligodendrocyte targets predominantly captured cell– cell adhesion and synapse-contact annotations (including presynaptic membrane assembly/organization and neurexin-associated pathways), reflecting conserved axon–glia interface signaling modules rather than broad lineage programs per se.

Finally, we evaluated enhancer-level predictions against experimentally validated BICCN Armamentarium enhancers. Enhancers ranked highly by DeepSCENIC for cell-type–specific activity were enriched for validated enhancers and showed agreement between predicted and measured activity across cell types (Fig. 4f-g, Methods). Notably, enhancers with higher experimental cell-type specificity exhibited stronger concordance with DeepSCENIC predictions, consistent with the model capturing cell-type–specific regulatory programs.

Together, these analyses indicate that DeepSCENIC recovers conserved transcriptional regulators, enhancer sequence logic, and downstream regulatory connections across human and mouse cortex, providing cross-species support for inferred GRNs in the absence of comprehensive experimental ground truth.

## DISCUSSION

The development of computational strategies to decode genomic regulatory programs has largely followed two separate directions. On one side, sequence-to-function (S2F) models have achieved remarkable success in deciphering the cis-regulatory code, yet they remain largely blind to the trans-environment, or cellular context, that provides combinations of transcription factors, their dosage, and their co-factors. On the other side, GRN inference methods have modeled relationships between transcription factors (TFs) and genes, but have often ignored the rich combinatorial code and function of genomic enhancers, as well as the complex interactions between distal enhancers and target genes. Here, we unify these paradigms by merging S2F models with single-cell GRN inference through a new deep learning framework, DeepSCENIC. By making TF activity a function of both sequence-derived binding potential and cell-specific TF expression, DeepSCENIC brings the trans-environment into a sequence-aware architecture. This allows the model to predict chromatin accessibility and gene expression at single-cell resolution using the TF expression vector of each cell as input. Importantly, the DeepSCENIC architecture was designed for biological interpretability. Because the model is mechanistically parametrized to learn two distinct matrices, one representing TF-to-region binding and one representing region-to-gene regulation, the learned parameters are not abstract weights hidden in a black box, but are directly linked to biology, encoding the physical connectivity of the GRN.

DeepSCENIC sits at the intersection of three lines of work that have so far developed largely in isolation. The first comprises local cis-regulatory S2F models with small receptive fields, such as BPNet^2^, ChromBPNet^6^ and CREsted^5^, which model TF binding or chromatin accessibility at base-or peak-level resolution. These models excel at decoding the internal grammar of individual enhancers such as motif syntax, affinity, spacing and combinatorial co-occurrence, and support detailed interpretation through in silico mutagenesis and de novo motif discovery. However, these models carry no representation of the trans-environment: a learned sequence feature is never explicitly assigned to the TF protein that recognizes it. Furthermore, these models do not connect an enhancer to the gene it regulates. DeepSCENIC exploits the strengths of enhancer representations from this family of models. For example, our dS-CREsted variant uses a CREsted backbone, and our in silico sequence-evolution experiments recover canonical motifs de novo for GATA1, HNF4A and others. Yet, in DeepSCENIC, enhancers are embedded within an explicit TF–RE–TG network, so that sequence-derived regulatory features are linked to named TFs and propagated to downstream genes.

The second line of work consists of gene-locus models, or genomic foundation models, with large receptive fields such as Enformer^18^, Borzoi^19^, and most recently AlphaGenome^20^. These models are trained to predict chromatin and transcriptional readouts across hundreds of kilobases to a megabase and can, in principle, learn enhancer-to-gene relationships directly through attention. In practice, however, the effective range of these models is considerably shorter than their nominal input, and the contribution of distal enhancers is systematically underweighted relative to promoter-proximal sequence^21^. This limits their reliability precisely for the task of assigning distal regulatory elements to target genes that lies at the heart of GRN inference. DeepSCENIC takes a deliberately different route to long-range regulation: rather than requiring a single network to propagate signal across an entire locus, it decouples the problem. Local sequence grammar is modelled per ATAC peak using exactly the kind of S2F backbone described above, including Enformer and Borzoi themselves as feature extractors, while enhancer-to-gene coupling is captured by explicit, learnable RE–TG weights (β) over peaks within ±1 Mb of the TSS. This separation lets DeepSCENIC exploit the cis-grammar while side-stepping their distal-enhancer attribution problem. It also collapses the per-gene readout to a scalar expression value, a design choice shared with recent single-cell sequence models such as Decima^50^, which reduces each gene to a single predicted log(CPM+1) per pseudobulk rather than a full base-resolution coverage track.

The closest precedents to DeepSCENIC are recent context-aware S2F models that begin to incorporate the trans-environment into sequence-based prediction. Corgi^35^ conditions a long-range S2F model on quantitative TF expression, and Scooby^36^ couples Borzoi-derived sequence embeddings with single-cell multiome representations to predict cell-state-specific profiles. Both demonstrate that sequence models can be made sensitive to cellular context, and Scooby in particular shares our strategy of fine-tuning a Borzoi backbone. They differ from DeepSCENIC, however, in objective and interpretability: they are built as accurate predictors of expression or accessibility in context, but they do not explicitly parameterize a TF–region–target gene graph, do not assign sequence features to named TFs, and are not constructed as mechanistic simulators that propagate a defined perturbation through learned regulatory links. DeepSCENIC instead exposes the GRN as the model’s internal wiring (θ and β), so that the same parameters that drive prediction can be read out as biology and directly intervened upon.

The third line of work is GRN inference itself. State-of-the-art methods such as SCENIC+^30^, Pando^31^ and CellOracle^32^ assemble TF–enhancer–gene networks from single-cell multiome data, but rely on Position Weight Matrix (PWM) scanning and statistical heuristics to place TFs on enhancers, and on correlation or distance priors to link enhancers to genes. These methods disregard most of the sequence grammar that the S2F models above have shown to be informative. While high-quality motif databases are now available, PWM-based scanning or enrichment alone remains limited by high false-positive rates and by its inability to model the broader sequence context, spacing, affinity, and combinatorial grammar through which TFs regulate enhancers. The enhancer–gene component has itself become an active subfield: dedicated methods such as scE2G^51^ link peaks to genes with supervised classifiers trained on CRISPRi perturbation data, combining ABC-style accessibility-and-distance features with peak–gene correlation across single cells, and achieve strong performance against CRISPR, eQTL and GWAS benchmarks. Such approaches are powerful, but they treat the enhancer as a feature vector rather than as a sequence, and thus cannot reason about which TFs bind or how a variant would alter that binding. The primary innovation of DeepSCENIC is to unify these strands: by leveraging transfer learning from S2F models like Enformer and Borzoi, it predicts TF–RE interactions from sequence at nucleotide resolution and learns RE–TG links jointly within the same objective, moving beyond simple motifs. Our results show that DeepSCENIC de novo recovers high-fidelity TF binding sites, as seen in our sequence evolution experiments for GATA1 and HNF4A, without being provided with prior motif knowledge for TFs, and that its enhancer–gene predictions, benchmarked against the same class of K562 CRISPRi data used to train scE2G, improve significantly over correlation-based baselines.

One of the most significant challenges in single-cell biology is predicting how a cell will respond to a perturbation, and dynamic, perturbation-aware modelling is itself not new to GRN inference. Both SCENIC+^30^ and CellOracle^32^ already simulate cell-state changes in response to TF perturbation: SCENIC+ fits random-forest regressors over a static GRN topology, while CellOracle propagates a TF knockout through a regularized linear GRN to estimate shifts in cell identity over a low-dimensional state manifold. In both cases, however, the simulator is a statistical model layered on top of a fixed network whose edges were placed using PWMs and accessibility–expression correlation, and the perturbation can only be expressed in the trans dimension, at the level of TF abundance. DeepSCENIC differs in two respects. First, its simulator is the network: perturbations propagate through the same sequence-grounded TF–RE–TG parameters that generated the predictions, rather than through a separate downstream regressor. The high concordance between in silico SOX10 knockdown and experimental data demonstrates that the model has captured the actual regulatory logic of the MEL-to-MES phenotype switch, in a manner that can be interpreted mechanistically. Second, because TF–RE activity is itself a function of the underlying DNA, this predictive power extends to the nucleotide level: DeepSCENIC not only predicted the impact of TF depletion but also the functional consequences of single-nucleotide variants in MEL-specific enhancers, remaining on par with specialized CNN models like DeepMEL. To our knowledge, this is the first model with the dual capability of simulating both trans-(TF levels) and cis-(DNA sequence) perturbations, including the ability of synthetic enhancer design at the cis-level, and the ability to perform genome-wide screens of TF knock-outs, TF over-expression, or combinations of TF perturbations.

A complementary and rapidly growing class of models approaches perturbation from the opposite direction. Virtual-cell and perturbation-foundation models, exemplified by the Arc Institute’s STATE^52^, are trained on hundreds of millions of observed and perturbed single cells. These models learn to map a starting transcriptome and a perturbation label directly to a perturbed expression state, through a learned latent space and large-scale supervision on perturbation atlases. These models are powerful precisely because they make no commitment to a mechanism: they capture perturbation responses statistically, without representing enhancers, motifs, or the cis-regulatory code at all. This comes at the cost of interpretability, generalizability, and of any handle on the DNA sequence, and therefore on genetic variation between individuals or species. DeepSCENIC is a smaller model that does not train on perturbational data, but is explicitly wired to the genome sequence.

While DeepSCENIC represents a step forward, it is not without limitations. We observed that the model struggled to fully disentangle TFs with highly overlapping binding profiles and expression patterns, such as HNF1A and HNF4A in HepG2 cells. This highlights a need for even higher-resolution data, perhaps through the integration of small-molecule perturbations or time-series single-cell data, to better distinguish between cooperative binding partners. Furthermore, while our RE-TG (enhancer-to-gene) predictions outperformed correlation-based baselines, they still trail behind fully supervised models trained on CRISPRi data. Also, our current implementation is by design limited to TF perturbations. It would be interesting in future developments to include protein-protein interaction networks into the model, to allow for non-TF perturbations such as signaling pathways.

DeepSCENIC thus provides a unified framework for understanding how the genome and the cell-specific trans-environment interact to produce a cell state. By integrating genomic foundation models into the heart of GRN inference, we have created a tool that is both biologically interpretable and predictive. We foresee that sequence-grounded GRN simulators and data-driven virtual-cell models could converge toward the same goal of a virtual cell model^53^ capable of predicting the effects of both mutations and drugs across the landscape of human cell types in health and disease, including the mechanistic, cis-regulatory foundation that explains how a perturbation produces the response it does.

## METHODS

### Model parameterization and outputs

DeepSCENIC is explicitly parameterized as an interpretable TF-RE-TG network, where the learnable parameters encode *trans-*(TF-RE) and *cis-*(RE-TG) regulatory effects. The model integrates (i) sequence-derived TF binding potentials *θ*, computed from a finetuned sequence-to-function (S2F) backbone and (ii) cell-specific TF expression from scRNA-seq to infer enhancer activities, which in turn predict both chromatin accessibility and target gene expression.

#### Notation

Let *t* ∈ {1, …, *T*} index TFs, *r* ∈ {1, …, *R*} index regulatory elements (REs), *g* ∈ {1, …, *G*} index genes (TGs), and *c* ∈ {1, …, *C*} index cells. For each region *r*, the S2F backbone processes the one-hot encoded DNA sequence and produces a TF-specific binding potential *θ_t_*_,*r*_. For each cell *c*, scRNA-seq provides a TF expression vector *x_t_*_,*c*_.

#### TF activity on regulatory elements (TF-RE)

DeepSCENIC models the activity *a_t_*_,*r*,*c*_ of TF *t* on region *r* in cell *c* as a linear combination of latent TF expression and sequence-derived binding potential:

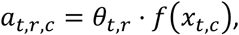

where each TF expression value *x_t_*_,*c*_ is passed through a shared non-linear variational encoder *f*(⋅) that operates on TFs independently. The encoder maps each scalar input *x_t_*_,*c*_ to a one-dimensional latent using a two-layer multilayer perceptron with tanh nonlinearities and a Gaussian variational output. To enforce non-negativity, all layers in the encoder use positively constrained weights by parametrizing the effective weight matrix as |*W*|, which enforces a monotone non-decreasing mapping from TF expression to latent expression (up to tanh saturation). During training, TF activities are sampled via the reparameterization trick:

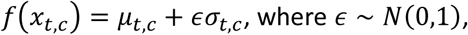

whereas during inference the posterior mean is used:

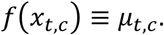

A Kullback-Leibler (KL) divergence term between the approximate posterior *N*(*μ_t,,c_*, σ^2^_*t,c*_) and a standard normal prior is added to the learning objective to regularize the TF expression latent.

#### Enhancer activity

The aggregate activity of a regulatory element is modeled as the total contribution of TF activities acting on this element:

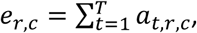

yielding a per-cell, per-enhancer latent representation that is constrained by both chromatin accessibility and gene expression during training.

#### Chromatin accessibility prediction

Chromatin accessibility is predicted directly from the inferred enhancer activities, thereby probing an explicit constraint on *e_r_*_,*c*_. For each cell *c*, the vector of enhancer activities *e<u>_c_</u>* = (*e*_1,*c*_, …, *e_R_*_,*c*_) is passed through a shared decoder ℎ(⋅) that output reconstructed accessibility for all enhancers:

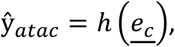

where ℎ(⋅) is a multilayer perceptron with positively constrained weights and tanh nonlinearities that maps each scalar *e_r_*_,*c*_ to a reconstructed chromatin accessibility value, using shared parameters across regions. Joint optimization of the ATAC reconstruction objective and the RNA reconstruction objective encourages enhancer activities *e_r_*_,*c*_ to capture regulatory variation that is simultaneously consistent with chromatin accessibility and downstream gene expression.

#### *Cis-*regulation to target genes (RE-TG)

Target gene expression is predicted as a weighted additive contribution of enhancer activities located within a ±1Mb window around the gene’s transcription start site. Let Ε(*g*) denote the set of REs in this window around gene *g*, and let *e_r_*_,*c*_ denote the inferred activity of RE *r* in cell *c*. DeepSCENIC learns RE-TG weights *β_r_*_,*g*_ and first compute a gene-specific regulatory input as:

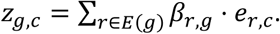

The predicted expression of gene *g* in cell *c* is obtained by passing this scalar through a shared non-linear decoder *q*(⋅) that operates on genes independently:

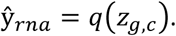

In our implementation, *q*(⋅) is a multilayer perceptron with tanh nonlinearities and positively constrained weights, shared across all genes. Thus, gene-specific behavior arises from the learned *cis-*regulatory weights *β_r_*_,*g*_ and enhancer activities *e_r_*_,*c*_, while *q*(⋅) provides a common non-linear mapping from aggregated regulatory input to the reconstructed expression scale.

#### Training objective and optimization

DeepSCENIC is trained end-to-end by jointly optimizing sequence-derived TF-RE binding potentials and the multiome recontraction of chromatin accessibility and gene expression. Training proceeds in two stages: (i) an initialization stage in which TF-RE binding potentials are computed for all regulatory elements, and (ii) a fine-tuning stage in which TF-RE binding potentials are refined while training the downstream TF-RE-TG network parameters to reconstruct observed single-cell profiles. To make fine-tuning scalable to large numbers of regulatory elements, sequence-derived TF-RE scores *θ* are updated stochastically for a subset of regions at each optimization step, while a cached estimate is retained for regions not updated in that step.

The overall learning objective is a weighted sum of reconstruction and regularization terms. For each cell *c*, the model minimizes a gene expression reconstruction loss *L_rna_* and a chromatin accessibility reconstruction loss *L_atac_*. Unless stated otherwise, *L_rna_* is the mean absolute error (MAE), whereas *L_atac_* is a cosine-similarity loss defined as *L_atac_* = 1 − cos(*y_ata_*_c_ − ŷ*_ata_*_c_). A KL divergence term *L_KL_* regularizes the variational TF-expression latent towards a standard normal prior (weighted by *φ*). In addition, interpretability is promoted through sparsity-induced penalties on *θ* and *β* regulatory parameters: (i) an *l*_1_-style penalty (*R_TF_*_−*RE*_ = |*θ*|) on TF-RE scores *θ* (weighted by *α*) and (ii) a distance-weighted sparsity penalty on RE-TG links (*R_RE_*_−*TG*_ = |*β*|), where each candidate region-gene pair (*r*, *g*) is weighted by a distance prior *γ*(*d_r_*_,*g*_):

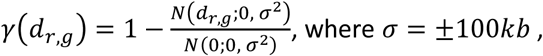

so that proximal links are penalized less than distal links. Overall, the objective can be summarized as:

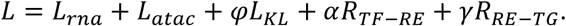

Model selection is performed on a held-out set by monitoring reconstruction performance, with the best checkpoint selected based on chromatin accessibility validation objective. To assess generalization across both cell states and genomic loci, we used a two-way split strategy: (i) a cell-wise split, in which a subset of cells is held out from training and used for test evaluation of both RNA and ATAC reconstruction; and (ii) a chromosome-wise split for chromatin accessibility, in which all regulatory elements (and corresponding sequences) from a subset of chromosomes are withheld for testing. This chromosome-wise split prevents leakage due to local genomic similarity and evaluates whether the learned sequence representations transfer to unseen loci.

#### Training protocol and hyperparameters

DeepSCENIC was trained end-to-end by jointly optimizing a sequence-to-function backbone, a TF–region prediction head, and a variational GRN module that reconstructs both gene expression and chromatin accessibility from single-cell multiome data. All S2F backbones were integrated into DeepSCENIC using an identical downstream architecture, loss function, and training schedule. Differences between model variants arise exclusively from the choice of pretrained sequence backbone and its associated sequence embedding (Table 1).

**Table 1.**
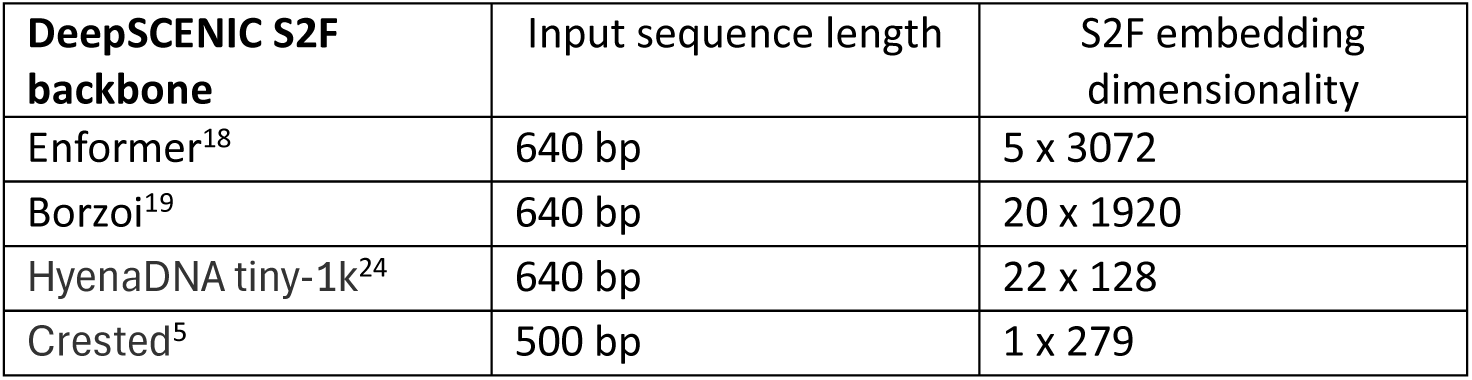
DeepSCENIC S2F backbone architectures. Overview of sequence-to-function (S2F) backbone models integrated into DeepSCENIC, including input sequence length and resulting embedding dimensionality.

The training pipeline consists of three coupled modules:

#### Sequence-to-function backbone

A pretrained genomic foundation model (Enformer, Borzoi, HyenaDNA, or CREsted, depending on the experiment) is used to extract sequence embeddings from fixed-length DNA sequences centered on scATAC-seq peaks. These embeddings are passed to a TF–RE prediction head that outputs TF-specific binding potentials (θ) from S2F sequence embeddings.

#### TF–RE head

The TF-RE head is a shallow neural network that maps S2F sequence embeddings to TF-specific binding scores for each regulatory element. These scores parameterize the TF–RE matrix (θ).

#### Variational GRN module (VAE)

The GRN module models TF activity as a variational latent variable inferred from TF expression, and sequence-derived TF binding potentials θ to compute enhancer activities, and predicts both chromatin accessibility and target gene expression through additive RE–TG effects (β).

Training proceeds in two stages. In the first stage, the sequence-to-function (S2F) backbone, the TF–RE head, and the variational GRN module were trained jointly on regulatory elements and genes from training chromosomes. Pretrained S2F weights were fine-tuned during this phase together with TF-RE head and variational GRN module parameters, minimizing the composite objective described above.

Optimization was performed using separate mini-batches for sequence and cellular components: regulatory element batches were used for backbone and TF–RE updates, whereas cell batches were used to optimize the GRN module. To maintain scalability with large peak sets, TF–RE binding potentials (θ) were updated stochastically: at each training step, only a subset of regulatory elements was forwarded through the S2F backbone and TF–RE head, while cached θ values were used for the remaining elements. Sequence batches were augmented using random positional shifts and reverse complements during training but not at test time.

Because RE–TG weights (β) are optimized jointly with θ under this partially cached update scheme, they are learned against a mixture of freshly updated and cached TF–RE scores. After convergence of the joint stage, we therefore performed a second optimization stage in which the S2F backbone and TF–RE head were frozen and TF–RE binding potentials were fully recomputed. The cis-regulatory RE–TG weights (β) were then re-optimized by freezing all other model parameters and using consistent, fixed θ values on training chromosomes and training cells. This retraining step ensures that enhancer–gene coupling is calibrated against the final sequence-derived TF–RE landscape rather than intermediate cached estimates.

In a final locus-generalization stage, regulatory elements located on held-out chromosomes were evaluated. Sequence-derived TF–RE scores for these regions were computed using the frozen S2F backbone, and corresponding RE–TG weights (β) were optimized using training cells and testing on held-out cells. This procedure assesses whether sequence representations learned on training loci generalize to unseen genomic regions and whether cis-regulatory coupling can be estimated for novel loci without further backbone adaptation.

## Extraction of cell-type–specific regulatory programs

Cell-type–specific regulatory programs were extracted from trained DeepSCENIC models by aggregating *trans-*and *cis-*regulatory weights across cells belonging to the same annotated cell type.

For each cell, TF–RE activity scores were computed as described in the “Model parameterization and outputs” section, combining sequence-derived TF binding potentials θ with the latent representation of TF expression. To obtain cell-type–specific TF–RE scores, TF–RE activities were averaged across all cells assigned to a given cell type, yielding a cell-type–level TF–RE activity matrix. This matrix quantifies, for each transcription factor and regulatory element, the predicted regulatory activity specific to that cell type.

*Cis-*regulatory RE–TG interactions were extracted from the learned RE–TG weight matrix β, which models enhancer–gene effects within the predefined genomic window. Because RE–TG weights are not cell-specific but are learned globally during training, cell-type specificity was introduced by weighting RE–TG interactions by cell-type–specific enhancer activity. Specifically, enhancer activities were averaged across cells of the same type, and RE–TG interactions were scaled to reflect the effective contribution of each regulatory element in that cellular context.

For each cell type, TF–target gene (TF–TG) scores were computed by composing the learned TF–RE binding potentials (θ) with the RE–TG regulatory weights (β), yielding a TF–gene coupling matrix (W) that reflects the structural *cis-*regulatory wiring:

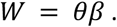

Cell-type specificity was then introduced by weighting each TF’s regulatory influence by its mean latent expression within that cell type:

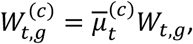

where μ̄^(*c*)^_*t*_ is the mean latent TF availability for TF *t* in cell type *c*.

Thus, TF–TG scores represent the effective regulatory influence of a TF in a given cellular context, integrating sequence-defined binding (θ), enhancer–gene coupling (β), and cell-type–specific TF availability.

To define cell-type master regulators, we first identified cell-type–specific regulatory elements using the inferred enhancer activity profiles. Specifically, for each cell type we performed differential testing on enhancer activities across cell types (Wilcoxon rank-sum), and retained the top K enhancers showing increased activity in that cell type (i.e., positively differentially active enhancers). We then quantified, for each TF, its effective average activity on the cell-type–specific enhancer set. TFs were finally ranked by the magnitude, and the top-ranked TFs were reported as cell-type master regulators, capturing TFs whose inferred availability aligns with strong sequence-based binding to enhancers that are specifically active in that cell type.

## In-silico perturbations

In-silico perturbations were performed by intervening on transcription factor (TF) expression and propagating the effect through the learned regulatory architecture without re-training the model. For a given perturbation, the expression of one or more TFs was set to a specified value (e.g., zero for knockdown or increased for overexpression) across all cells. The perturbed expression matrix was then forwarded through the variational encoder to obtain updated latent TF expression values. TF–RE activities were recomputed by combining the perturbed latent TF expression with the learned TF–RE binding potentials (θ), and enhancer activities were propagated through the fixed RE–TG weights (β) to obtain predicted gene expression changes. To account for indirect and higher-order effects mediated through the regulatory network, this procedure was iterated for a fixed number of steps, each time updating gene and TF expression. The final output of the perturbation simulation is a predicted perturbed gene expression matrix alongside log-fold changes relative to the unperturbed baseline, providing a mechanistically grounded estimate of the regulatory consequences of TF perturbation in a given cellular context.

For *cis-*regulatory perturbations, nucleotide-level modifications were introduced directly into regulatory element sequences. Mutated sequences were forwarded through the S2F backbone and TF-RE head to obtain updated TF–RE binding potentials (θ), while the *trans-*regulatory state (TF availability) was kept constant for the cell type of interest. The perturbed binding potentials were then propagated through the same enhancer–gene wiring (β) to compute updated TF–TG regulatory scores and predicted expression changes. Mutation effects were quantified as the difference between perturbed and wild-type regulatory influence, enabling mechanistic assessment of how sequence variation alters TF binding and downstream chromatin and gene regulation within a defined cellular context.

## Sequence evolution and motif recovery experiments

To assess whether DeepSCENIC captures coherent and TF-specific *cis-*regulatory sequence logic, we performed in-silico sequence evolution experiments using the trained S2F backbone and TF-RE head. Starting from either random DNA sequences or existing regulatory element sequences, we iteratively introduced single-nucleotide substitutions and evaluated their predicted TF–RE binding potentials *θ_t_*(*s*), where *s* denotes the input sequence and *θ_t_*(*s*) is the predicted binding potential for transcription factor *t*.

At each iteration, all possible single-base substitutions of the current sequence were generated and evaluated. For a chosen target TF *ṫ*, mutations were ranked according to a specificity-aware objective that promotes strong activation of the target TF while penalizing activation of competing TFs:

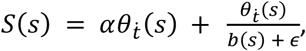

where *b*(*s*) denotes the maximum absolute prediction across off-target TFs, *α* rescales the activation term and *ε* ensures numerical stability. Both the target activation term and the specificity ratio were clipped at predefined upper bounds to prevent numerical instability and runaway optimization. At each step, the mutation that maximally increased this score was retained, yielding a greedy hill-climbing procedure. Optimization terminated when improvements fell below a predefined tolerance threshold or when the target activation exceeded a specified level while maintaining sufficient specificity.

For each TF, we performed 1,000 independent sequence evolution runs from independent random sequence initialization and analyzed the resulting evolved sequences jointly for motif recovery and robustness. Prior to motif analysis, evolved sequences were filtered to retain only high-confidence sequences exhibiting strong and specific activation of the target TF. Specifically, we retained sequences in which the predicted binding potential for the target TF exceeded a predefined activation threshold (>30) and in which the target prediction was at least as large as the strongest competing prediction among a predefined set of related TFs (specificity ratio ≥1). To recover sequence motifs from evolved sequences, we extracted high-contributing subsequences (i.e. seqlets) from evolved sequences by identifying contiguous regions that contributed most strongly to the target TF binding potential using in-silico mutagenesis profiles. Aggregating these subsequences across runs yielded position-specific sequence patterns, which were compared to known TF binding motifs. Successful recovery of canonical motifs indicates that the trained model encodes TF-specific *cis-*regulatory grammar.

## GRN benchmark evaluation protocol

### ChIP-seq recovery and correlation analysis

For each TF with available ENCODE ChIP-seq data, we evaluated concordance between predicted TF-RE binding potentials (θ) and experimental ChIP-seq signal measured over scATAC-seq consensus peak set. ChIP-seq coverage was quantified as normalized signal intensity across the consensus peaks. For each TF and each model variant, we computed the Pearson correlation coefficient between predicted TF-RE scores and ChIP-seq signal across all regions. This yielded one correlation value per TF per method. The distribution of per-TF correlation coefficients across TFs was used to compare methods.

To assess the ability of each method to prioritize experimentally supported binding regions, we performed a ranked recovery analysis. For each TF and each ChIP-seq track, consensus regions were ranked by ChIP-seq coverage and restricted to the top 10,000 regions. Independently, each model variant produced a TF-specific ranked list of predicted target regions, from which the top 10,000 regions were retained. We then computed a cumulative recovery curve along the ChIP-ranked list by counting, for each ChIP rank position, the cumulative number of ChIP regions that were also contained in the top 10,000 predicted regions. Curves were aggregated across TFs to compare model variants.

### Enhancer–target gene CRISPRi benchmark

To evaluate enhancer–gene (RE–TG) predictions, we benchmarked predicted *cis-*regulatory weights against experimentally validated enhancer–target gene pairs derived from ENCODE CRISPRi-based perturbation screens in K562 cells. The ground-truth dataset consists of enhancer–gene links identified through CRISPR interference experiments, in which candidate distal regulatory elements were individually perturbed and target gene expression responses were measured. Candidate RE–TG pairs were defined as all regulatory elements located within ±1 Mb of a gene’s transcription start site, consistent with the candidate window used during model training. For each gene, all candidate regulatory elements within this window were considered as potential positives or negatives. For each model variant, RE–TG pairs were ranked according to their predicted regulatory score (RE–TG weight). Experimentally validated RE–TG links were treated as positive examples, while non-validated candidate pairs within the same genomic window were treated as negatives. Performance was quantified using receiver operating characteristic (ROC) analysis, computing the area under the ROC curve (AUC) to measure how well validated RE–TG pairs were prioritized over non-validated candidates. As baselines, we ranked RE–TG pairs using correlation-based approaches, including Pearson correlation between enhancer accessibility and gene expression across cells, with and without distance-based weighting (i.e. applying the same distance prior used to regularize β weights), and evaluated them using the same ROC framework.

### TF perturbation benchmark

To evaluate inferred TF–target gene (TF–TG) relationships, we benchmarked predicted TF–TG scores against differential expression profiles from ENCODE TF knockout experiments. For each perturbed TF, genes were labeled as responders (“positives”) if they were significantly differentially expressed, defined as *p* < 0.01 and |log_2_ *FC*| > 0; all other assayed genes were treated as negatives. Genes were ranked by the absolute predicted TF–TG score, and performance was quantified using the area under the receiver operating characteristic curve (ROC AUC). AUC values were computed independently for each TF perturbation experiment and compared across methods.

### MM lines analysis

To evaluate the ability of DeepSCENIC to model phenotype switching in a disease-relevant context, we analyzed a melanoma single-cell multiomic atlas comprising melanocytic (MEL), intermediate (INT), and mesenchymal-like (MES) cell states.

### Definition of the MEL–MES transcriptional axis

A principal component analysis (PCA) was performed on log-normalized gene expression values across all melanoma cells. The first principal component (PC1) captured the dominant transcriptional continuum separating MEL and MES phenotypes and was therefore used as a quantitative MEL–MES axis.

For each cell c, the position along the phenotypic axis was defined as its PC1 coordinate. Perturbation-induced state transitions were quantified as:

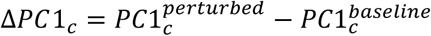

Positive and negative shifts correspond to movement towards MES and MEL programs, respectively.

Genome-wide in silico TF screen along the MEL–MES axis. To systematically identify regulators of phenotype switching, each TF was individually knocked down in silico. TF knockdowns were simulated by setting the expression of a selected TF to zero across all cells and propagating the perturbation through the inferred DeepSCENIC gene regulatory network for a fixed number of iterative update steps (N = 5).

The resulting perturbed expression matrix was projected onto the MEL–MES transcriptional axis (PC1), and transcription factors were ranked by the magnitude and direction of the induced axis shift within each state, thereby enabling identification of regulators driving transitions toward MES or MEL programs.

### Validation against experimental SOX10 knockdown

To assess predictive accuracy, simulated SOX10 knockdown effects were compared to experimental time-series RNA-seq data following SOX10 depletion in intermediate melanoma lines. Concordance analysis was restricted to experimentally responsive genes in order to focus on biologically meaningful perturbation effects. To this end, differentially expressed genes were defined as those with if *p_adj_* < 0.05 and |log_2_ *F C*| > 2.

Concordance between predicted and observed fold changes was quantified using Pearson correlation and Concordance Correlation Coefficient (CCC).

In addition, MEL and MES program trajectories were quantified using the curated Verfaillie program genes^45^. For each intermediate line, we restricted the comparison to Verfaillie genes that were shared between the DeepSCENIC simulation and the SOX10 knockdown time course and that showed a minimal predicted response (mean |log2FC| > 0.05 across iterations). For each retained gene, we computed the Pearson correlation between its predicted log2FC trajectory across simulation iterations (iter1–iter4) and its experimentally measured expression changes across the SOX10 knockdown time points (NTC/BL, 24 h, 48 h, 72 h). The resulting per-gene correlations were summarized per line to compare predicted and experimental dynamics.

### Quantitative validation of enhancer activity upon sequence perturbation

To quantitatively evaluate predictive accuracy at single-nucleotide resolution, we compared DeepSCENIC-predicted mutation effects with experimentally measured enhancer activity changes for three melanoma-specific synthetic enhancers. For each enhancer, single-nucleotide variants were introduced in silico, and predicted regulatory effects were computed as log2 fold changes in enhancer-driven gene expression relative to the wild-type sequence. These predictions were directly compared to experimentally measured MPRA log2 fold changes obtained from luciferase assays for the corresponding mutations.

### Sequence-level explanation of the IRF4 enhancer

To interpret and quantitatively validate sequence determinants underlying state-specific enhancer activity, we performed in silico mutagenesis (ISM) on a melanoma-active IRF4 enhancer. Single-nucleotide substitutions were systematically introduced across the enhancer sequence, and the resulting changes in predicted enhancer activity were quantified using the trained DeepSCENIC model.

For comparison to experimental measurements, we included MPRA-based in vitro mutagenesis (IVM) data from Kircher et al. in the SK-MEL-28 cell line, which provide direct measurements of activity changes for individual nucleotide substitutions. As model-based baselines, we computed DeepLIFT attribution scores from the melanoma-specific convolutional model DeepMEL, and gradient × input attribution scores from Enformer (interpreted for the SK-MEL-5 class). DeepSCENIC mutagenesis profiles were computed separately for MEL, INT, and MES cellular contexts.

### Cross-species GRN conservation analysis

Cross-species comparisons were restricted to one-to-one human–mouse orthologs obtained from Ensembl (release 115). All conservation analyses were performed independently per matched cortical cell type unless otherwise stated.

#### TF activity conservation

For each species, TF activity scores were inferred per cell type using DeepSCENIC. Within each matched cell type, orthologous TFs were ranked by activity separately in mouse and human. Overlap between top-ranked TF sets was quantified using Jaccard similarity for varying thresholds. Statistical significance was assessed by comparing observed overlaps to a null distribution generated by random sampling of TFs from the shared ortholog universe (2,000 permutations per k). Normalized enrichment scores (NES) were computed as:

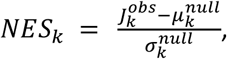

where *J^obs^_k_* denotes the observed Jaccard similarity between the top-k TF sets in mouse and human, and *μ^null^_k_* and *σ^null^_k_* represent the mean and standard deviation of the Jaccard similarities obtained under random sampling at the same k.

#### Enhancer regulatory code conservation

To assess conservation of *cis-*regulatory sequence logic, we compared mouse and human regulatory regions based on their sequence-derived TF binding potential profiles inferred by DeepSCENIC human model. For each species, enhancers were represented by vectors of predicted TF activities. Mouse and human enhancer matrices were concatenated and projected into a shared low-dimensional space using principal component analysis (100 principal components). A k-nearest neighbor (kNN) graph (k=15) was then constructed across the combined enhancer set using cosine similarity in principal component space. Conservation was quantified using a neighborhood enrichment framework: for each enhancer, we computed the fraction of nearest neighbors belonging to the same annotated cell type but originating from the opposite species. Observed cross-species same-cell-type neighbor fractions were compared to random expectation obtained by permuting cell-type labels within species (1,000 permutations). Normalized enrichment score over random expectation was used as a measure of conserved enhancer regulatory code.

#### TF–target gene conservation

Species-specific TF–target gene (TF–TG) networks were constructed by integrating TF–RE links RE–TG links. For each TF, target genes were ranked by aggregated TF–TG score separately in mouse and human. Conservation was assessed independently per TF by comparing the top-k predicted targets (k = 200) across species using Jaccard similarity. Significance was evaluated using a gene-level random sampling null model over the shared orthologous gene universe, and NES values were calculated as described above.

#### Evaluation against BICCN Armamentarium enhancers

To evaluate enhancer-level predictions, we compared DeepSCENIC-predicted enhancer activities to experimentally measured enhancer activity from the BICCN Armamentarium AAV reporter assays^48^. For each enhancer, predicted activity was computed as the mean inferred enhancer activity across cells within each annotated cortical cell type and min-max normalized per enhancer. Experimental activity values (log CPM) were similarly aggregated per enhancer and cell type. For every enhancer, we computed the Pearson correlation coefficient between predicted and experimentally measured activity profiles across matched cell types. Statistical significance was determined using two-sided Pearson correlation tests.

To assess whether prediction accuracy depended on enhancer specificity, experimental enhancer specificity was quantified as the maximum normalized activity fraction across cell types. Enhancers were binned into six equal-width bins according to their correlation coefficients, and mean enhancer specificity (± SEM) was computed per bin. The relationship between prediction accuracy and enhancer specificity was visualized by plotting per-enhancer correlation values, colored by maximum specificity, alongside the binned summary statistics.

In addition, we evaluated cell-type–specific prediction performance as a binary classification task. For each cell type, enhancers were labeled as active in each cell type based on experimental reporter signal (log CPM > 10). Predicted enhancer specificity scores were used as classifier scores, and performance was quantified using area under the receiver operating characteristic curve (AUROC) and area under the precision–recall curve (AUPR) for each cell type.

## Datasets

### ENCODE deeply profiled cell lines

To benchmark the accuracy of GRN inference with DeepSCENIC, we used simulated single-cell multiome data from eight deeply profiled ENCODE cell lines, as originally introduced in the SCENIC+ study^30^. Bulk RNA-seq and ATAC-seq were downloaded from the ENCODE portal for MCF7 (ENCFF136ANW; ENCFF772EFK), HepG2 (ENCFF660EXG; ENCFF239RGZ), PC3 (ENCFF874CFD; ENCFF516GDK), GM12878 (ENCFF626GVO; ENCFF415FEC), K562 (ENCFF833WFD; ENCFF512VEZ), Panc1 (ENCFF602HCV; ENCFF836WDC), IMR90 (ENCFF027FUC; ENCFF848XMR), and HCT116 (ENCFF766TYC; ENCFF724QHH). For each cell line, 500 simulated single-cell profiles were generated for each cell line by randomly sampling 50,000 RNA-seq reads and 20,000 ATAC-seq fragments from their corresponding bulk transcriptomic and chromatin accessibility profiles, resulting in a total of 4,000 paired single-cell multiomic profiles. Gene expression data were further filtered to retain genes expressed in at least 10 cells, then normalized using total-count scaling followed by log-transformation yielding 15,276 genes, including 1,594 TFs used as inputs to the model.

Chromatin accessibility was summarized over a consensus peak set of fixed width (500 bp; 558,315 peaks in total). Importantly, DeepSCENIC was not trained on raw ATAC fragment counts, instead, we used topic-based denoised peak accessibility inferred with cisTopic^54^ as the chromatin input. All sequences and genomic coordinates were based on the human reference genome hg38.

To evaluate generalization across both cell states and genomic loci, we used a two-way split strategy. First, we performed a cell-wise split holding out 20% of cells for test evaluation of both RNA and ATAC reconstruction (80/20 train/test). Second, to assess sequence-level generalization, we held out all peaks from chromosomes chr19, chr18, chr11, chr7, and chr3 as an out-of-chromosome test set (430,589 training regions; 127,726 test regions).

Finally, to define candidate *cis-*regulatory links, we constructed a region to gene association matrix by assigning each peak to all genes with transcription start sites within ±1Mb, which defines the candidate RE-TG search space used by the *cis-*regulatory module.

### Melanoma cell lines

To evaluate DeepSCENIC in a disease relevant setting with well characterized phenotype switching, we used the melanoma cell line multiomic dataset from the SCENIC+ study^30^. This resource comprises nine patient derived melanoma clusters spanning melanocytic (MEL), intermediate (INT), and mesenchymal-like (MES) transcriptional states. Processed scRNA-seq count matrix and corresponding scATAC-seq topic-denoised imputed accessibility matrix were downloaded from SCENIC+ resources. Gene expression was filtered and log-normalized, yielding 12169 genes, including 1386 transcription factors used as DeepSCENIC input. CisTopic imputed accessibility peaks were used as chromatin accessibility input to the model.

To assess generalization across both cell states and genomic loci, we used the same two-way split strategy as above. First, we performed 80/20% train/test split over the full dataset of 936 paired profiles for evaluation of RNA and ATAC reconstruction. Second, to evaluate sequence-level generalization, we performed an out-of-chromosome split by holding out all peaks on chromosomes chr16, chr18, and chr19 (259,169 training peaks; 22,652 test peaks). Finally, candidate *cis-*regulatory links were defined by assigning each peak to all genes with transcription start sites within ±1 Mb (hg38), which defines the RE– TG search space used by the *cis-*regulatory module.

### Human and mouse motor cortex multiome atlases (BICCN/BICAN)

To study conservation of GRNs across species in a complex in vivo tissue, we used single-cell multiome atlases of the primary motor cortex (M1) from the BICCN/BICAN consortium^47^, which provide matched scRNA-seq and scATAC-seq profiles across major cortical subclasses.

#### Human cortex

The human dataset comprised 31,960 cells with paired scRNA-seq and scATAC-seq profiles. Gene expression was filtered to retain genes detected in at least 10 cells, then log-normalized, resulting in a set of 16,735 genes, of which 1,588 TFs used as DeepSCENIC input. For chromatin accessibility, we used imputed accessibility values provided after topic modelling, and we restricted the peak set to regions supported within each cortical subclass to reduce extremely sparse/rare peaks. For each subclass, we retained peaks detected in at least 1% of cells (449 filtered peaks). The final evaluation used the same two-way split strategy as above: an 80/20 cell-wise train/test split and an out-of-chromosome split holding out all peaks on chr19, chr18, chr11, and chr7 (509,775 training regions; 102,935 test regions).

#### Mouse cortex

The mouse dataset comprised 39,470 cells with paired scRNA-seq (16,437 genes) and scATAC-seq profiles (540,627 peaks). RNA preprocessing mirrored the human dataset (min 10 cells per gene; total-count normalization; log-transformation). For ATAC, we analogously used cisTopic-derived peak accessibility values and applied a stricter subclass-support filter to focus on robust regulatory elements across mouse subclasses: within each subclass we retained peaks detected in at least 10% of cells and took the union across subclasses (2,344 filtered peaks). We applied the same two-way evaluation design (80/20 cell-wise split; chr19, chr18, chr11, chr7 held out for out-of-chromosome testing), resulting in 438,728 training regions and 101,899 test regions, with 1,390 TFs as model inputs.

Candidate *cis-*regulatory links for both species were defined by assigning each peak to all genes with transcription start sites within ±1 Mb (hg38), providing the candidate RE–TG search space for the *cis-*regulatory module.

## Code availability

The source code for the DeepSCENIC package is available at https://github.com/aertslab/deepSCENIC. Code and notebooks used to reproduce the analyses, benchmarking experiments and figures in this manuscript are provided in a separate repository https://github.com/aertslab/deepSCENIC_analyses.

## Data availability

This study did not generate new primary experimental data requiring deposition. All datasets used in this study are publicly available from the original publications or repositories cited in the manuscript. Accession numbers and links for the single-cell multiome, chromatin accessibility, gene expression, and sequence-to-function model backbones are provided in Supplementary Table 1. For the TF–RE and TF-TG benchmarks, ChIP-seq datasets, bulk RNA-seq experiments upon TF perturbation were obtained from the SCENIC+ publication^30^. Processed input matrices, trained models and intermediate files required to reproduce the main figures and results are available at DOI: https://doi.org/10.48804/FPJDLP.

## Author contributions

G.P. and S.A. conceived the study. G.P. developed the DeepSCENIC framework, performed the computational analyses, and implemented the modeling and benchmarking experiments. S.D.W. contributed to data preprocessing. C.B. and V.K. assisted with integration of sequence-to-function backbone models. G.P. and S.A. wrote the manuscript with input from all authors. S.A. supervised the study.

## Acknowledgements

We thank members of the Aerts laboratory and the VIB.AI Center for AI & Computational Biology for insightful discussions and feedback. We acknowledge the Vlaamse Supercomputer Center and VIB datacore, and technical support teams for providing high-performance computing resources used in this study. This research was funded in part by FWO (G0I2722N EOS ID 40007513, G094121N and G044124N); Foundation Against Cancer (2020-1396 and 2024-140); Inter-university BOF-projects (IBOF/25/01); ERC AdG (101054387 Genome2Cells).

## References

1. Zhou, J. & Troyanskaya, O.G. Nat Methods 12, 931–934 (2015).

2. Avsec, Ž., et al. Nat Genet 53, 354–366 (2021).

3. Kelley, D.R., Snoek, J. & Rinn, J.L. Genome Res. 26, 990–999 (2016).

4. Yuan, H. & Kelley, D.R. 2021.09.08.459495 (2021).doi:10.1101/2021.09.08.459495

5 Kempynck, N. et al.2025.04.02.646812 (2025).doi:10.1101/2025.04.02.646812

6. Pampari, A. et al.2024.12.25.630221 (2025).doi:10.1101/2024.12.25.630221

7. Sundararajan, M., Taly, A. & Yan, Q. (2017).doi:10.48550/arXiv.1703.01365

8. Shrikumar, A., Greenside, P. & Kundaje, A. (2019).doi:10.48550/arXiv.1704.02685

9. Shrikumar, A., et al. (2020).doi:10.48550/arXiv.1811.00416

10. Winter, S.D. et al.2026.01.14.699402 (2026).doi:10.64898/2026.01.14.699402

11. Janssens, J. et al. Nature 601, 630–636 (2022).

12. Bravo González-Blas, C., et al. Nat Cell Biol 26, 153–167 (2024).

13. Hecker, N. et al. Science 387, eadp3957 (2025).

14. Minnoye, L. et al. Genome Res. 30, 1815–1834 (2020).

15. Sarropoulos, I. et al. Science 391, eadw9154 (2026).

16. Taskiran, I.I. et al. Nature 626, 212–220 (2024).

17. Winter, S.D. et al.2026.01.14.699402 (2026).doi:10.64898/2026.01.14.699402

18. Avsec, Ž., et al. Nat Methods 18, 1196–1203 (2021).

19. Linder, J., Srivastava, D., Yuan, H., Agarwal, V. & Kelley, D.R. Nat Genet 57, 949–961 (2025).

20. Avsec, Ž., et al. Nature 649, 1206–1218 (2026).

21. Karollus, A., Mauermeier, T. & Gagneur, J. Genome Biol 24, 56 (2023).

22. He, A.Y., Palamuttam, N.P. & Danko, C.G. 2024.10.15.618510 (2025).doi:10.1101/2024.10.15.618510

23. Sasse, A. et al. Nat Genet 55, 2060–2064 (2023).

24. Nguyen, E., et al. (2023).doi:10.48550/arXiv.2306.15794

25. Sanabria, M., Hirsch, J., Joubert, P.M. & Poetsch, A.R. Nat Mach Intell 6, 911–923 (2024).

26. Dalla-Torre, H. et al. Nat Methods 22, 287–297 (2025).

27. Zhou, Z., et al. (2024).doi:10.48550/arXiv.2306.15006

28. Brixi, G. et al.2025.02.18.638918 (2025).doi:10.1101/2025.02.18.638918

29. Tomaz da Silva, P., et al. Nat Genet 57, 2589–2602 (2025).

30. Bravo González-Blas, C., et al. Nat Methods 20, 1355–1367 (2023).

31. Fleck, J.S. et al. Nature 621, 365–372 (2023).

32. Kamimoto, K. et al. Nature 614, 742–751 (2023).

33. Yuan, Q. & Duren, Z. Nat Biotechnol 43, 247–257 (2025).

34. Saraswat, M. et al.2025.05.13.653733 (2025).doi:10.1101/2025.05.13.653733

35. Aksu, E.D. & Vingron, M.2025.06.25.661447 (2025).doi:10.1101/2025.06.25.661447

36. Hingerl, J.C. et al. Nat Methods 22, 2275–2285 (2025).

37. Smith, R.P. et al. Nat Genet 45, 1021–1028 (2013).

38. Yu, M. et al. Molecular Cell 36, 682–695 (2009).

39. Wingelhofer, B. et al. Leukemia 32, 1713–1726 (2018).

40. Mui, A.L., Wakao, H., Kinoshita, T., Kitamura, T. & Miyajima, A. EMBO J 15, 2425–2433 (1996).

41. Gupta, S., Stamatoyannopoulos, J.A., Bailey, T.L. & Noble, W.S. Genome Biol 8, R24 (2007).

42. Winter, S.D. et al.2026.01.14.699402 (2026).doi:10.64898/2026.01.14.699402

43. Gschwind, A.R. et al.2023.11.09.563812 (2023).doi:10.1101/2023.11.09.563812

44. Wouters, J. et al. Nat Cell Biol 22, 986–998 (2020).

45. Verfaillie, A. et al. Nat Commun 6, 6683 (2015).

46. Kircher, M. et al. Nat Commun 10, 3583 (2019).

47. Callaway, E.M. et al. Nature 598, 86–102 (2021).

48. Ben-Simon, Y. et al. Cell 188, 3045–3064.e23 (2025).

49. Stergachis, A.B. et al. Nature 515, 365–370 (2014).

50. Lal, A. et al. (2024).doi:10.1101/2024.10.09.617507

51. Sheth, M.U. et al. (2024).doi:10.1101/2024.11.23.624931

52. Adduri, A.K. et al.

53. Bunne, C. et al. Cell 187, 7045–7063 (2024).

54. Bravo González-Blas, C., et al. Nat Methods 16, 397–400 (2019).

