## Supplementary figures and images for "DeepSCENIC: transfer learning from sequence-to-function models enables causal gene regulatory network inference"

### Supplementary Figure 1

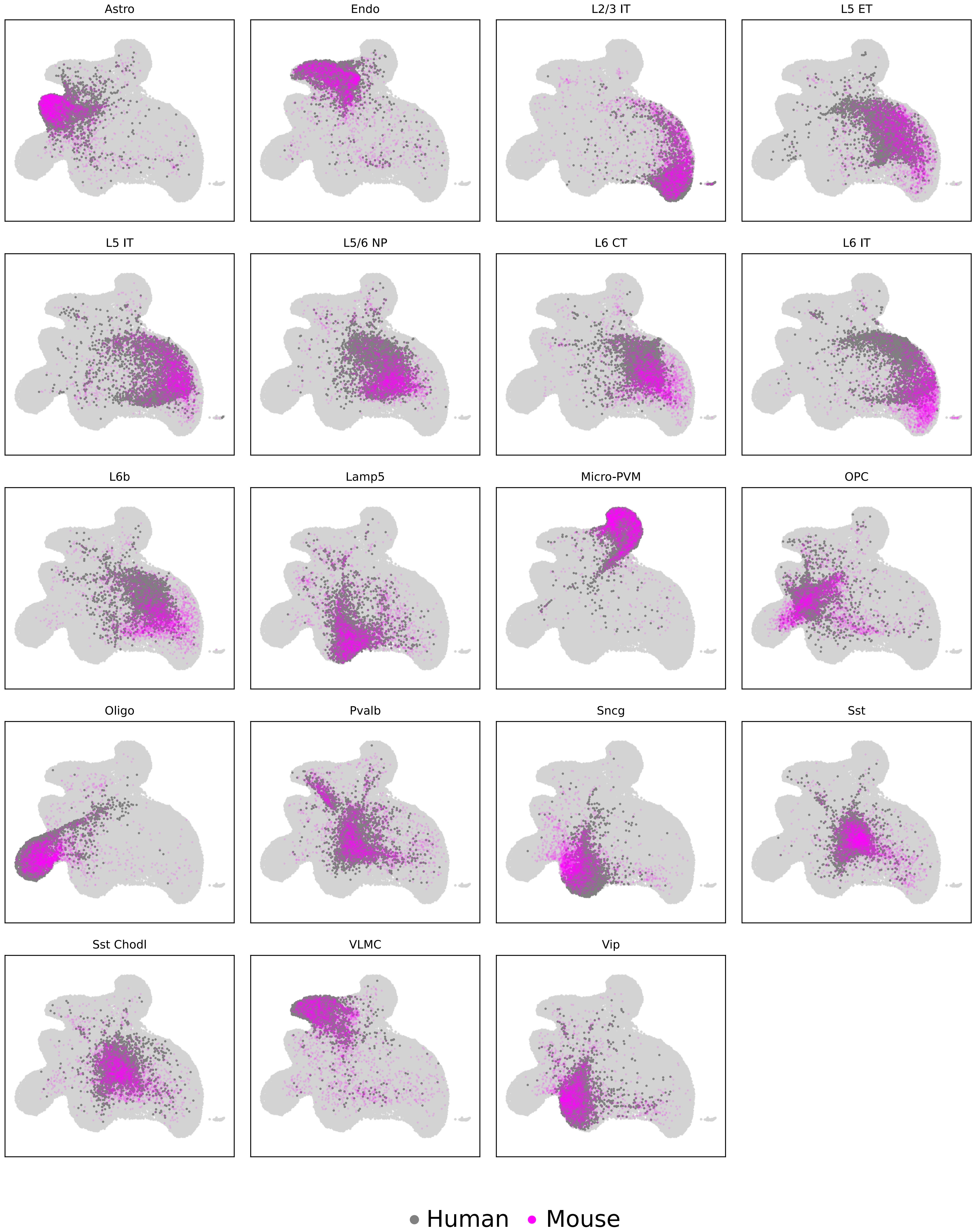
